# Elucidation of protein-protein interactions necessary for maintenance of the BCR-ABL signaling complex

**DOI:** 10.1101/684480

**Authors:** Tomas Gregor, Michaela Kunova Bosakova, Alexandru Nita, Sara P. Abraham, Bohumil Fafilek, Nicole H. Cernohorsky, Jan Rynes, Silvie Foldynova-Trantirkova, Jiri Mayer, Lukas Trantirek, Pavel Krejci

## Abstract

Approximately 50% of chronic myeloid leukemia (CML) patients in deep remission experience a return of clinical CML after withdrawal of tyrosine kinase inhibitors (TKIs). This suggests signaling of inactive BCR-ABL, which allows for survival of cancer cells, leading to relapse. Understanding the dynamics of BCR-ABL signaling complex holds a key to the mechanism of BCR-ABL signaling. Here, we demonstrate that TKIs inhibit catalytic activity of BCR-ABL, but do not dissolve the BCR-ABL core signaling complex consisting of CrkL, SHC1, Grb2, SOS1, cCbl, and SHIP2. We show that CrkL binds to proline-rich regions located in C-terminal, intrinsically disordered region of BCR-ABL, that deletion of pleckstrin homology domain of BCR-ABL diminishes interaction with SHC1, and that BCR-ABL sequence motif located in disordered region around phosphorylated tyrosine 177 mediates binding of at least three core complex members, the Grb2, SOS1 and cCbl. Introduction of Y177F substitution blocks association with Grb2, SOS1 and cCbl. Further, we identified SHIP2 binding sites within the src-homology and tyrosine kinase domains of BCR-ABL. We found that BCR-ABL is unable to phosphorylate SHC1 in cells lacking SHIP2. Reintroducing SHIP2 into Ship2 knock-out cells restored SHC1 phosphorylation, which depended on inositol phosphatase activity of SHIP2. Our findings provide characterization of protein-protein interactions in the BCR-ABL signaling complex, and support the concept of targeting BCR-ABL signaling in CML by inhibition of its interactions with the members of the core complex.

## Introduction

Chronic myeloid leukemia (CML) is a clonal myeloproliferative disorder characterized by the t(9;22)(q34;q11) translocation. This translocation generates a fusion oncogene containing part of the breakpoint cluster region (*BCR*) gene joined with *ABL* gene, which encodes a tyrosine kinase. The resulting protein has a constitutive tyrosine kinase activity caused by oligomerization of the BCR domains, which promote autophosphorylation-mediated activation of ABL kinase domain (Zhao et al. 2002). The tyrosine kinase activity of BCR-ABL is necessary and sufficient to maintain the CML phenotype, and acquisition of active BCR-ABL generates a CML-like lethal leukemia in mice models, both confirming the central role of BCR-ABL in the pathogenesis of CML (Voncken et al. 1995). CML initially manifests as chronic leukemia, caused by expansion of myeloid lineage, but eventually accelerates into the blastic phase, characterised by massive increase of undifferentiated CML progenitors. CML is a lethal when untreated or in case of a failed treatment. At the cellular level, BCR-ABL signalling transforms hematopoietic cells by increasing proliferation and survival, and decreasing dependency on extracellular signals delivered by cytokines and growth factors (Hazlehurst et al. 2009). This is mediated by constitutive, BCR-ABL-mediated activation of large amount of signaling intermediates, including members of Ras/Erk MAP kinase pathway involved in cell proliferation, PI3K/Akt signaling conferring resistance to apoptosis, and Jak/STAT signaling contributing to cytokine independence (Skorski et al. 1997; Shuai et al. 1996; Steelman et al. 2004).

Suppression of BCR-ABL tyrosine kinase activity with small molecule tyrosine kinase inhibitors (TKI) have greatly improved CML prognosis. Imatinib was the first TKI approved to target BCR-ABL, and represented a major therapeutic breakthrough (Druker et al. 1996). Over a decade of clinical experience with imatinib demonstrated an estimated 85% of survival rate for the first-line treatment patients. When complete cytogenetic response is not achieved in first year of imatinib administration, the likelihood of CML progression or loss of imatinib response is 38%. This, together with the worse prognosis for patients incompletely responding to imatinib, leaves about third of CML patients with potential for improvements over the imatinib therapy (Cilloni and Saglio 2012). Failed imatinib response often involves *BCR-ABL* gene amplification, increased expression or occurrence of mutations causing imatinib resistance (Gorre et al. 2001; Modugno 2014). This was overcome by development of second generation of TKIs, nilotinib and dasatinib, which inhibit BCR-ABL with greater efficiency than imatinib, and target the majority of imatinib-resistant BCR-ABL mutants (Weisberg et al. 2005; Shah et al. 2005). Nilotinib and dasatinib provide a significant improvement in CML treatment over imatinib, inducing 2-year complete cytogenetics response in ∼40% of imatinib-resistant patients (Hochhaus et al. 2008). Unfortunately, in many CML patients who failed to respond to imatinib, a T315I substitution in the BCR-ABL’s kinase domain occurs. T315I targets the gate-keeper residue controlling access to the hydrophobic cavity adjacent to the ATP binding site, which is important for the proper drug binding (T. Zhou et al. 2007). The resistance BCR-ABL-T315I to TKIs was successfully addressed by development of ponatinib, which induces complete cytogenetic response in 46% of patients resistant to both nilotinib and dasatinib (Cortes et al. 2013).

In summary, although a direct targeting of BCR-ABL tyrosine kinase activity by TKIs generates impressive results in CML treatment, it fails primarily in three areas. First, some CML patients who harbour unmutated BCR-ABL remain resistant to TKIs, suggesting that other oncogenic pathways cooperate with BCR-ABL or that BCR-ABL tyrosine kinase activity is not necessary for CML persistence in these patients (Bewry et al. 2008). Second, resistance to TKIs eventually develops in significant percentage of CML patients. Novel mutations in BCR-ABL, resistant even to ponatinib, have been described, including the dual mutations affecting one BCR-ABL allele (Eide et al. 2011). It is expected that successive therapy with different TKIs will lead to continual selection of novel mutations in BCR-ABL (Modugno 2014).

Finally, TKIs suppress but not eradicate CML. The slowly proliferating leukemia stem cells are poorly targeted with TKIs (Corbin et al. 2011). The heterogenous cell pool that constitutes CML originates in a rare population of stem cells capable to recapitulate CML even after the deep molecular remission. This is evident in TKI discontinuation trials, which report a return of clinical CML after TKI withdrawal in approximately 50% of CML patients in deep remission, suggesting that long-term blockade of BCR-ABL kinase activity by TKIs alone is not sufficient to cure CML (Rousselot et al. 2014). This, together with other adverse aspects of TKI use such as their side effects and high economic costs of the life-long therapy necessitates development of conceptually novel treatments for CML. To achieve this goal, we first need to completely understand the mechanics of BCR-ABL signal transduction, as the BCR-ABL may play other roles beyond the constitutively active tyrosine kinase. The protein-protein interactions within the BCR-ABL signaling complexes may remain preserved when its kinase activity is inhibited by TKIs, leading to residual signaling necessary for survival of CML cells.

The main downstream signaling pathways utilized by BCR-ABL to regulate cell functions are well established. In contrast, the composition of BCR-ABL interactome, i.e. the pool of signaling intermediates associating directly with BCR-ABL is only beginning to emerge. Active BCR-ABL is phosphorylated on tyrosines 177, 1127 and 1294 (Mitra et al. 2013), which serve as docking sites for binding of proteins containing SH3 and PTB domains. Several such proteins have been identified, including adapters Gab2, CrkL and SHC1, adapter/phosphatase SHP2, p85 subunit of the PI3-kinase, ubiquitin ligase cCbl, and others (Brehme et al. 2009). This study was carried-out to map the protein-protein interactions within the BCR-ABL signaling complex in detail, and elucidate the dynamics of the BCR-ABL signaling complex in the active and TKI-inhibited state of BCR-ABL.

## Results and discussion

### Inhibition of BCR-ABL kinase activity does not dissolve the BCR-ABL signaling complex

TKIs inhibit kinase activity of BCR-ABL but may not interfere with the protein-protein interactions within the BCR-ABL signaling complex, particularly those which are not mediated by phosphorylated tyrosine motifs. We asked whether the inhibition of BCR-ABL kinase activity results in disintegration of its signaling complex. We expressed p210 BCR-ABL in 293T cells, inhibited its catalytic activity by nilotinib, and compared the size of BCR-ABL complexes in active and inactive state by ultracentrifugation in 15-40% sucrose gradient. Cell treatment by 100 nM nilotinib lead to complete suppression of BCR-ABL kinase activity, evidenced by the lack of autophosphorylation at Tyr412 (Fig. 1A). Inhibition of BCR-ABL kinase activity resulted in a partial shift in the BCR-ABL complexes towards lighter sucrose fractions, suggesting partial dissociation of the BCR-ABL signaling complex. The members of BCR-ABL core signaling complex p85α-PI3K, GRB2, SHIP2, SHC1, SOS1, SHP2 and cCBL (Brehme et al. 2009) co-sedimented with BCR-ABL in sucrose gradient (Fig. 1A); virtually no co-sedimentation was found with CRK, CRKL or GAB2 (not shown). Despite the complete inhibition of the BCR-ABL activity, evidenced as the lack of autophosphorylation at Tyr412, nilotinib did not cause the exclusion of any of the endogenously expressed interactors from the co-sedimentation with BCR-ABL. Quantification of western blot analysis of proteins saturated on BCR-ABL, i.e. those which majority co-sedimented with BCR-ABL in 293T cells (p85α-PI3K, GRB2 and SHIP2) shows that portion of GRB2, but not SHIP2 or p85α-PI3K dissociated from the BCR-ABL complex after nilotinib treatment (Fig. 1B), suggesting only partial dissolution of BCR-ABL signaling complex in 293T cells treated with nilotinib. Similar data were found in 293T cells transfected with p210 BCR-ABL and in K562 cells, a permanent cell line established from CML patient, which express endogenous BCR-ABL (Figs. S1, S2). The association of BCR-ABL with GRB2 and SHIP2 in 293T cells expressing p190 BCR-ABL was confirmed by proximity ligation assay (PLA). In PLA analyses, inhibition of BCR-ABL kinase activity by nilotinib lead to statistically-significant decrease in interaction with co-transfected GRB2 or SHIP2, but at least 50% of the interaction was preserved for both partners, when compared to active BCR-ABL (Fig. 1C). Next, the protein lysates of NIH3T3 cells expressing p210 BCR-ABL were resolved by blue-native (BN)-PAGE to separate protein complexes, which were then analyzed by second-dimension SDS-PAGE to obtain their individual components (Fig. 2A, B). Immunoblotting revealed ∼ 600-kDa protein complex containing BCR-ABL, SHIP2 and GRB2 (Fig. 2C). Quantification of the percentage of bound GRB2 and SHIP2 shows that inhibition of BCR-ABL catalytic activity with nilotinib or kinase inactivating mutations K271H (Preyer, Vigneri, and Wang 2011) did not inhibit SHIP2 association with BCR-ABL, while it significantly suppressed GRB2 association (Fig. 2D). Yet an approximately 30% of GRB2 still associated with kinase-inactive BCR-ABL, or BCR-ABL with Y177F substitution, which is known to disable the GRB2 binding motif on BCR-ABL (Pendergast et al. 1993; Goga et al. 1995).

**Figure 1.**
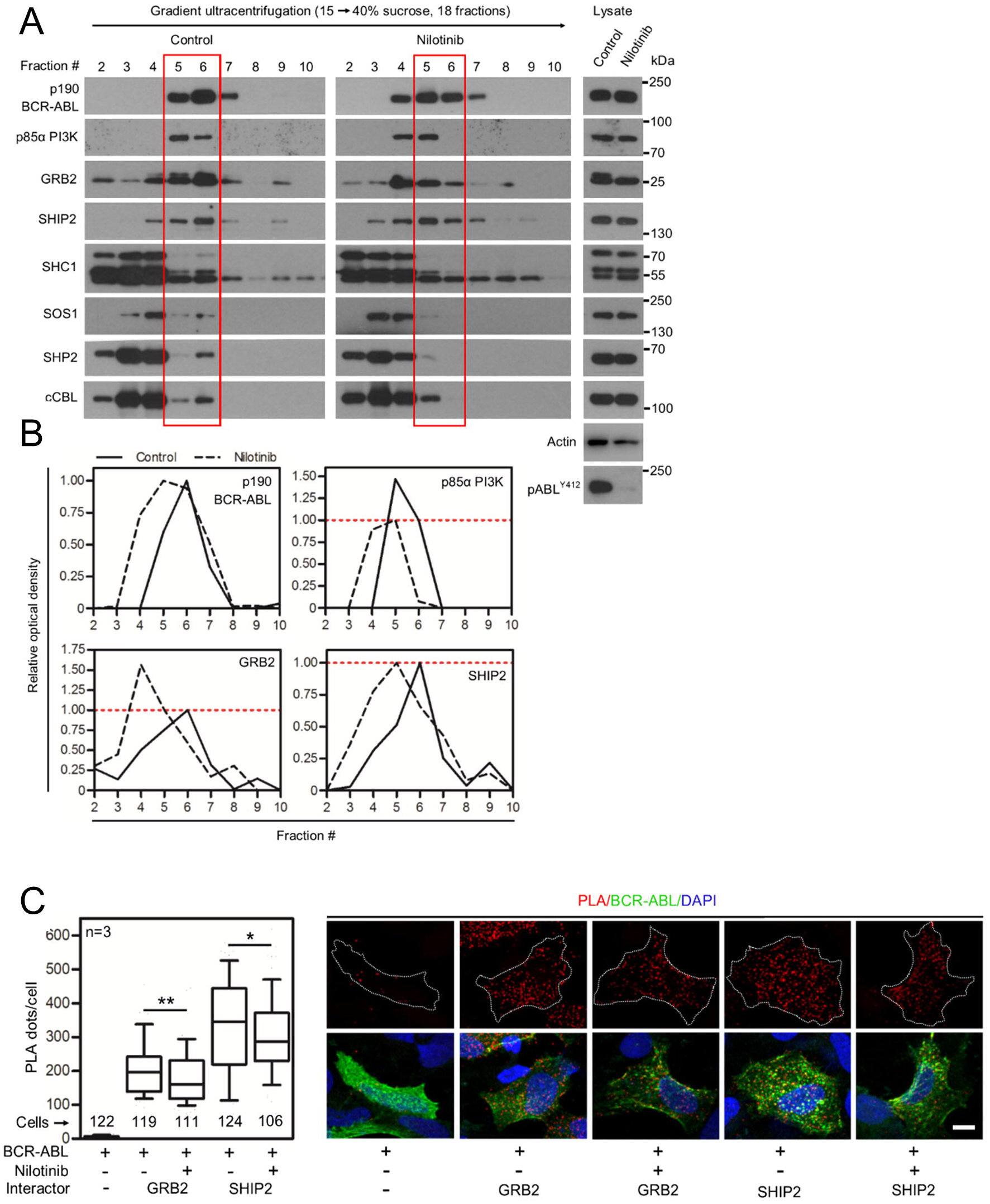
Nilotinib causes only partial dissolution of the BCR-ABL signaling complex. (A) 293T cells were transfected with p190 BCR-ABL, native cells lysates were subjected to ultracentrifugation in the 15-40% sucrose gradient, and the collected fractions were analyzed by western blot. Note the various degree of co-sedimentation of BCR-ABL with p85α PI3K, GRB2, SHIP2, SHC1, SOS1, SHP2 and cCBL. Inhibition of BCR-ABL kinase activity with 100 nM nilotinib resulted in a shift of a fraction of the BCR-ABL complexes towards lighter fractions, suggesting partial dissolution of the BCR-ABL signaling complex. (B) The western blot analysis of proteins co-sedimenting with BCR-ABL (p85a-PI3K, GRB2 and SHIP2) was quantified as described in Material and methods. Note that portion of GRB2, but not SHIP2 or p85a-PI3K dissociated from the BCR-ABL complex after nilotinib treatment. Data represent a single experiment out of three independent experiments carried-out. The fractions containing most of the p190 BCR-ABL are highlighted in red. Phosphorylation (p) at ABL Y412 was used to determine the degree of BCR-ABL inhibition using nilotinib; actin serves as a loading control in total cells lysates used for ultracentrifugation. (C) Cells were transfected with FLAG-tagged p190 BCR-ABL, V5-tagged GRB2 or SHIP2, and treated with nilotinib, and subjected to PLA. The antibodies against protein tags were used in PLA (red); cABL antibody was used to counterstain the transfected cells (green). Cells transfected with BCR-ABL and an empty vector serves as negative control. Number of PLA dots per cell was calculated and graphed (10-90 percentile). Statistically dignificant differences were highlighted (Student’s *t*-test with Welch’s correction for unequal variances; \**p*<0.05, \*\**p*<0.01). Scale bars, 10 µm.

**Figure 2.**
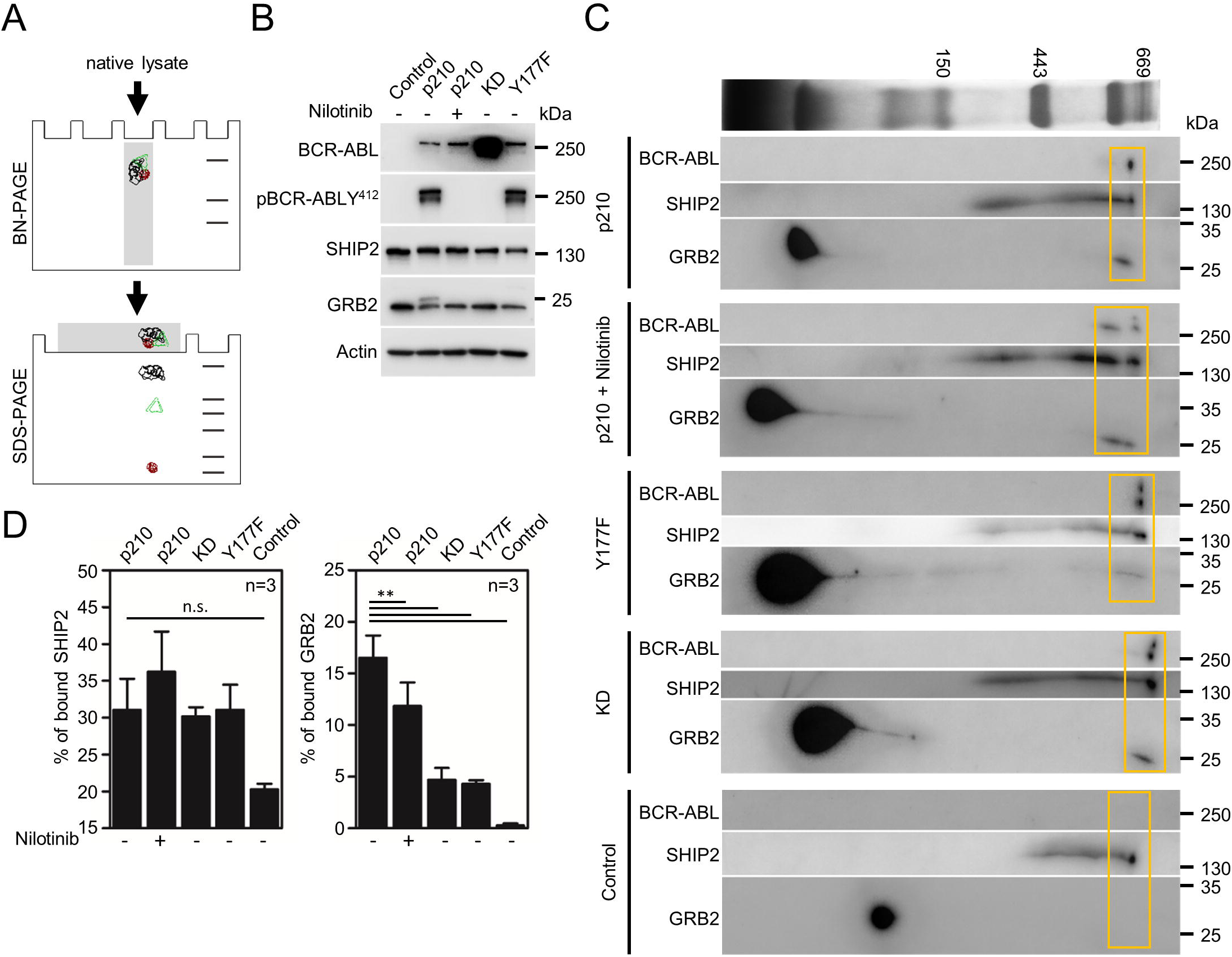
SHIP2 and GRB2 associate with catalytically inactive BCR-ABL. (A) Scheme of the experimental procedure comprising native lysis, blue native (BN)-PAGE, SDS-PAGE and western blot. The protein complexes are highlighted in color. (B) Cell lysates of NIH3T3 cells transfected with p210 BCR-ABL. KD, kinase-dead BCR-ABL mutant; Y177, BCR-ABL Y177F mutant. The inhibition of BCR-ABL kinase activity by nilotinib is demonstrated by lack of phosphorylation (p) at ABL residue Y412. Actin serves as loading control. (C) Merged second dimension BN-PAGE blots of cells transfected with different BCR-ABL variants. The membranes have been probed sequentially for BCR-ABL, SHIP2 and GRB2, the BCR-ABL/SHIP2/GRB2 complexes are highlighted by yellow box. (D) The quantification of the percentage of bound GRB2 and SHIP2 to the BCR-ABL. Statistically significant differences are highlighted (Student’s t-test, \*\**p*<0.01; n.s., not significant). Data are representative for three independent experiments.

Finally, the immunoprecipitation was used to study the integrity of BCR-ABL complex. 293T cells were transfected by p190 or p210 variants of BCR-ABL and BCR-ABL association with endogenously expressed SOS1, SHIP2, cCBL, SHC1 and p85a-PI3K was probed by co-immunoprecipitation. All studied interactors co-immunoprecipitated with both variants of BCR-ABL (Fig. 3A). Nilotinib reduced this association, but significant amounts of SOS1, SHIP2, cCBL and SHC1 still co-immunoprecipitated with BCR-ABL; no association of p85a-PI3K with BCR-ABL was found in cells treated with nilotinib (Fig. 3A, green arrows). Because endogenous STS1 and CRKL were not detected by western blot in 293T cells, transient transfection followed by immunoprecipitation was used to probe their interaction with BCR-ABL. Nilotinib suppressed, but did not abrogate STS1 and CRKL interaction with BCR-ABL (Fig. 3B, C; green arrows); similar results were obtained in experiments probing STS1 and CRKL association with BCR-ABL KD mutant (Fig. 3B, C; blue arrows). The endogenous GRB2 is difficult to detect in BCR-ABL immunocomplexes because is co-migrates with IgL. We therefore transfected GRB2 into 293T cells and probed association of BCR-ABL with GRB2 immunocomplexes. Figure 3D shows significant association of GRB2 with BCR-ABL, kinase-inactive due to the nilotinib treatment or KD mutations.

**Figure 3.**
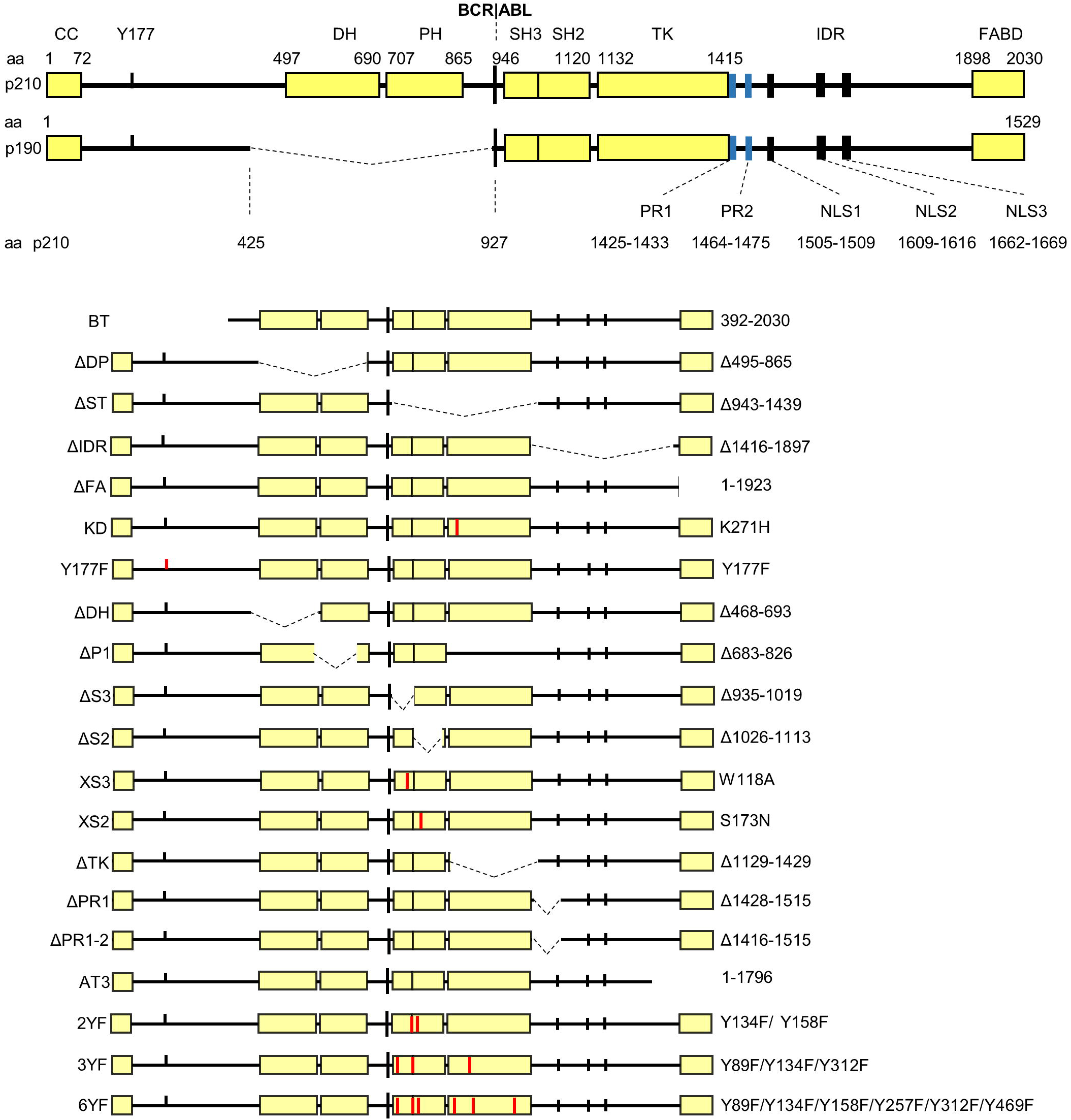
The BCR-ABL signaling complex is preserved after nilotinib treatment. 293T cells were transfected with p190 and p210 BCR-ABL alone (A) or together with STS1 (B), CRKL (C) and GRB2 (D). Cells treated with nilotinib (150 nM) are indicated by green arrows. Blue arrows indicate transfection with BCR-ABL variant devoid of kinase activity (KD), as evidenced by autophosphorylation (p) at ABL residue Y412. BCR-ABL was immunoprecipitated (IP) and binding of interaction partners was analyzed by western blot. Nilotinib diminished, but not completely inhibited binding of most interactors to BCR-ABL, with the exception of p85a-PI3K, which did not immunoprecipitate with BCR-ABL upon nilotinib treatment. Data are representative for three independents experiments (n=3). Actin serves as loading control in total cell lysates used for immunoprecipitation.

### Interaction of GRB2, SOS1, cCBL and SHC1 with BCR-ABL

Secondary structure prediction of p210 BCR-ABL indicates two disordered regions, located between the CC and DH domains, and between TK and FABD domains (Figs. 4, 5A). Because the peptide microarrays may be used to elucidate protein-protein interaction epitopes in both structured and unstructured proteins, we used this technology to characterize, in detail, the binding epitopes between BCR-ABL and members of its core complex. Thirteen amino acid long peptides corresponding to the primary sequence of p210 BCR-ABL were spotted on microarrays, incubated with interacting protein of interest, and analyzed as described in Material and Methods (Fig. 5B).

**Figure 4.**
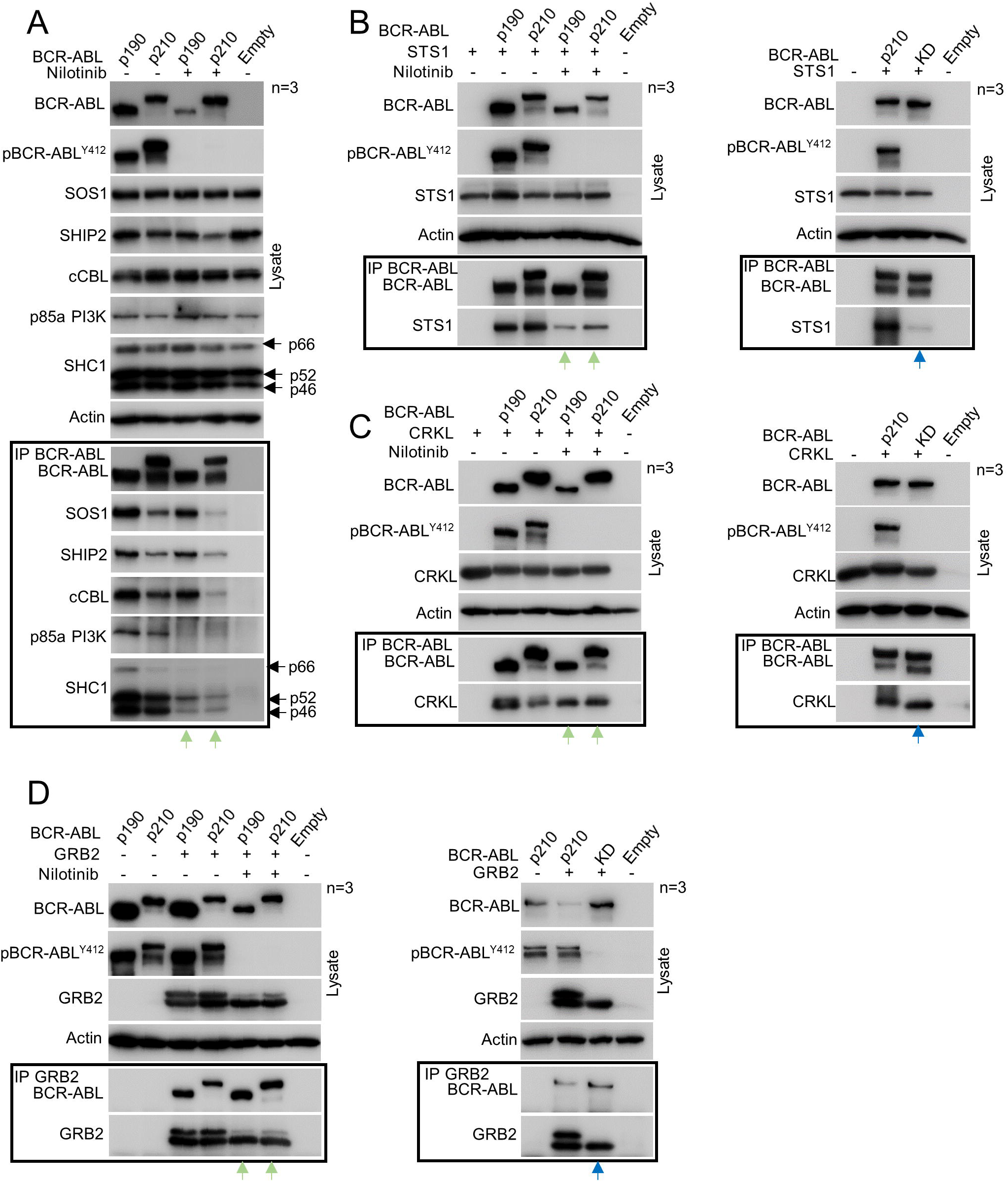
BCR-ABL variants used in the study. Schematic representation of generated p190 and p210 BCR-ABL variants. All constructs contain N-terminal FLAG epitope. The amino acid substitution are indicated in red; CC, coiled coil domain; DH, double homology domain; PH, pleckstrin homology domain; SH2, SH3, Src homology domain 2 and 3; TK, tyrosine kinase domain; IDR, intrinsically disordered region; FABD, F-actin binding domain; NLS, nuclear localization signal; PR, proline rich domain.

**Figure 5.**
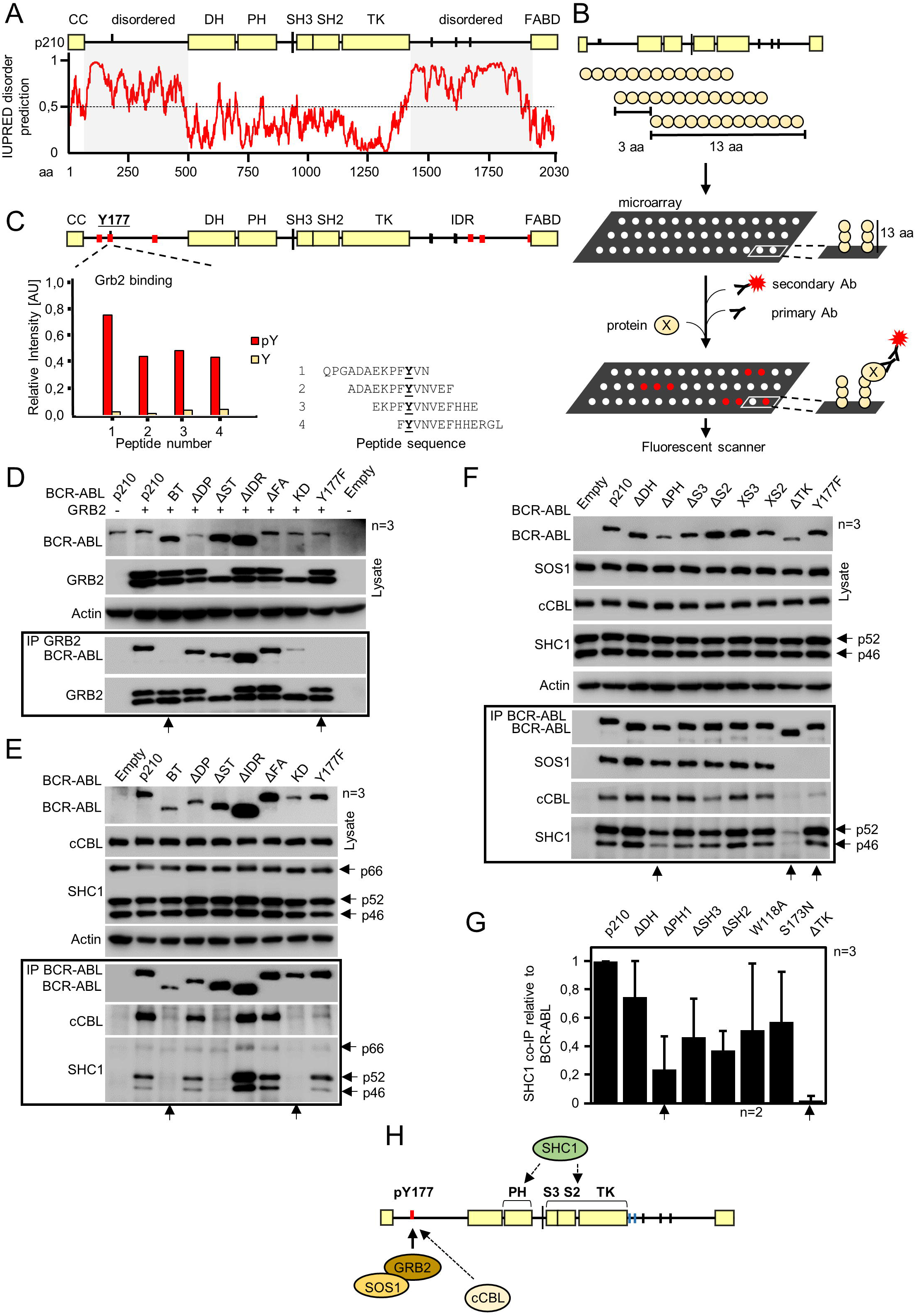
Interaction of GRB2, SOS1, cCBL and SHC1 with BCR-ABL. (A) Secondary structure prediction of p210 BCR-ABL by IUPRED. Values above 0.5 indicate two disordered regions on the BCR-ABL N- and C-termini, involving Y177 and three NLS, respectively. (B) Scheme of the microarray analysis. Thirteen amino acid long peptides corresponding to the primary sequence of p210 BCR-ABL were spotted on microarrays, incubated with protein of interest, primary and fluorescently labeled secondary antibodies, and scanned. Fluorescence intensity values for each spot were used to indicate binding of protein to BCR-ABL peptides. (C) Microarrays indicate direct binding of GRB2 to phosphorylated Y177. Red lines on BCR-ABL scheme indicate potential binding sites. Graph shows intensities for phosphorylated (red) and non-phosphorylated peptides involving peptides with Y177. (D) Immunoprecipitation (IP) of BCR-ABL with GRB2 after expression in 293T cells; Y177F substitution abrogates Grb2 association with BCR-ABL, similar to kinase-dead BCR-ABL (arrows). (E, F) Co-immunoprecipitation of endogenous cCBL, SHC1 and SOS1 with transfected BCR-ABL in 293T cells. Y177F abrogates binding of SOS1 and largely limits binding of cCBL (arrows). (H) Scheme of the proposed interaction. Grb2 binds directly to phosphorylated Y177 and recruits SOS1. pCbl requires pY177 and proline rich (PR) motif for binding. SHC1 requires TK domain and also partially requires pleckstrin homology (PH) domain for binding. Data are representative for three independent experiments (n=3).

Microarray data indicated that GRB2 binds to Y177 in BCR-ABL, and that this interaction is highly specific for phosphorylated version of Y177; the same Y177 peptides lacking the phosphorylation showed no interaction with GRB2 (Fig. 5C). Furthermore, microarrays revealed five more potential binding sites, two of them in disordered region of BCR and three in disordered region of ABL (Fig. 5A, Fig. S1A). Three of these sites contained PxxxR motif that is partial consensus binding motif of Grb2 C-terminal SH3 domain (PxxxRxxKP) (Lewitzky et al. 2001). To verify these binding sites, we generated several deletion variants of p210 BCR-ABL (Fig. 4), and probed the co-immunoprecipitation of these variants with V5-tagged GRB2 in 293T cells. BCR-ABL mutant Y177F had completely abolished binding to GRB2, as well as BCR-ABL with deleted N-terminal part of BCR including Y177 (BCR-ABL-BT) (Fig. 5D). Interaction of KD-BCR-ABL with GRB2 was largely limited, but still detectable. Given that Y177 is known to be autophosphorylated by BCR-ABL, it is possible that in KD BCR-ABL, other proteins are able to phosphorylate Y177 and create GRB2 binding site, as found before (Warmuth et al. 1997; Meyn et al. 2006). Deletion of IDR did not have effect on Grb2 binding and GRB2 did not bind to construct expressing only IDR sequence, suggesting these sites are not essential for GRB2 binding (Fig. 5H and data not shown).

Microarray data yielded no potential binding sites for cCBL despite extensive optimization and use of two different recombinant cCBL proteins (Fig. S1B). Interestingly, co-immunoprecipitations of endogenous cCBL revealed that Y177F substitution almost entirely abolished cCBL interaction with BCR-ABL. Because cCBL is known constitutive GRB2 interactor in various cells types (Buday et al. 1996; Donovan et al. 1994; Meisner et al. 1995; Panchamoorthy et al. 1996), we speculate that cCBL binds indirectly to BCR-ABL via GRB2 (Fig. 5H). This explains loss of cCBL binding to Y177F mutant that is defective for GRB2 binding. Furthermore, inactivation of BCR-ABL kinase activity (i.e. the ability to auto-phosphorylate Y177), by deletion of SH3-SH2-TK and TK domains, or introduction of KD substitutions also lead to loss of cCBL binding (Fig. 5E, F).

SHC1 is an adaptor protein that is capable of phosphotyrosine binding and it has three isoforms, p66, p52 and p46 which contain N-terminal PTB domain and C-terminal SH2 domain that flank central collagen homology 1 region. The PTB domain recognizes minimal NPXpY motif and extended LXNPTpY motif is needed for high affinity interaction (Trb et al. 1995). These motifs are not present in BCR-ABL. Microarray data showed multitude of potential binding sites mainly focused into the IDR region; no phosphotyrosine binding sites were identified (Fig. S1C, Tab. S4). SHC1 binding depended on tyrosine kinase activity of BCR-ABL, as the KD-BCR-ABL, ΔSH3-SH2-TK-BCR-ABL and BT constructs did not bind SHC1 (Fig. 5E). Similarly, deletion of TK domain again almost entirely abolished SHC1 binding (Fig. 5F, arrow). Deletion of PH domain also showed decreased interaction with SHC1 by approximately 70% (Fig. 5F, 5G). To evaluate SHC1 binding more precisely we used individual deletions of SH3, SH2 and TK domains, as well as substitutions W118A and S173N, inactivating proline rich binding function of SH3 and phosphotyrosine binding function of SH2, respectively (Brasher, Roumiantsev, and Van Etten 2001; Grebien et al. 2011). These individual deletions as well as substitutions decreased SHC1 interaction, however only deletion of TK domain abolished interaction almost completely (Fig. 5F, 5G).

### Interaction of CRKL and SHIP2 with BCR-ABL

CRKL adapter is a major substrate of BCR-ABL that is heavily phosphorylated in CML cells (Oda et al. 1994; ten Hoeve et al. 1994; Nichols et al. 1994). CRKL was shown to bind proline rich motifs APELPTKTR (PR1) and EPAVSPLLPRK (PR2) in IDR motif of ABL (R. Ren, Ye, and Baltimore 1994). The contribution of these sites for CRKL interaction with BCR-ABL however remains controversial, as the removal of entire motifs compromises binding, while more subtle deletions show no effect (Senechal, Halpern, and Sawyers 1996; Sattler et al. 2002; Heaney et al. 1997). Our microarray data showed four potential binding sites in structured regions of BCR-ABL, including phosphopeptides with tyrosines Y89 in SH3 domain, Y134 in SH3-SH2 linker and Y312 in TK domain and a peptide within FABD region (Fig. 6A). Peptides corresponding to the PR1 and PR2 sites were also positive, but were not formally considered as binding site as we only take into account three or more consecutive peptides with positive signal. The co-immunoprecipitation experiments showed that deletion of IDR domain eliminated binding to CRKL (Fig. 6B). Deletion of SH3-SH2-TK regions also abolished interaction with CRKL, however this deletion included also the PR1 site. Co-IP with BCR-ABL variants ΔPR1 and ΔPR1-2 containing smaller deletions within the IDR region showed that deletion that encompasses PR1 and PR2 sites has the biggest effect on CRKL binding (Fig. 6B; arrows). The individual deletion of SH2 and SH3 domains has no effect on binding, in contrast to deletion of TK domain showed reduction in CRKL binding, however this deletion also encompassed PR1 site. To evaluate if phosphorylated tyrosines Y89, Y134 and Y312 contribute to the CRKL interaction with BCR-ABL, we exchanged all three tyrosines for phenylalanines (BCR-ABL-3YF). This mutant interacted with CRKL normally, suggesting that these signals were false positives. Similarly, the interaction with FABD, identified by microarray, was not confirmed by co-immunoprecipitation (Fig. 6D). As cCBL is known CRKL interactor, we evaluated binding of cCBL to BCR-ABL in CRKL co-immunoprecipitation experiments as well. cCBL interaction with ΔPR1 and ΔPR1-2 mutants is limited as well, suggesting that some fraction of cellular cCBL binds to BCR-ABL via CRKL (Fig. 7B).

**Figure 6.**
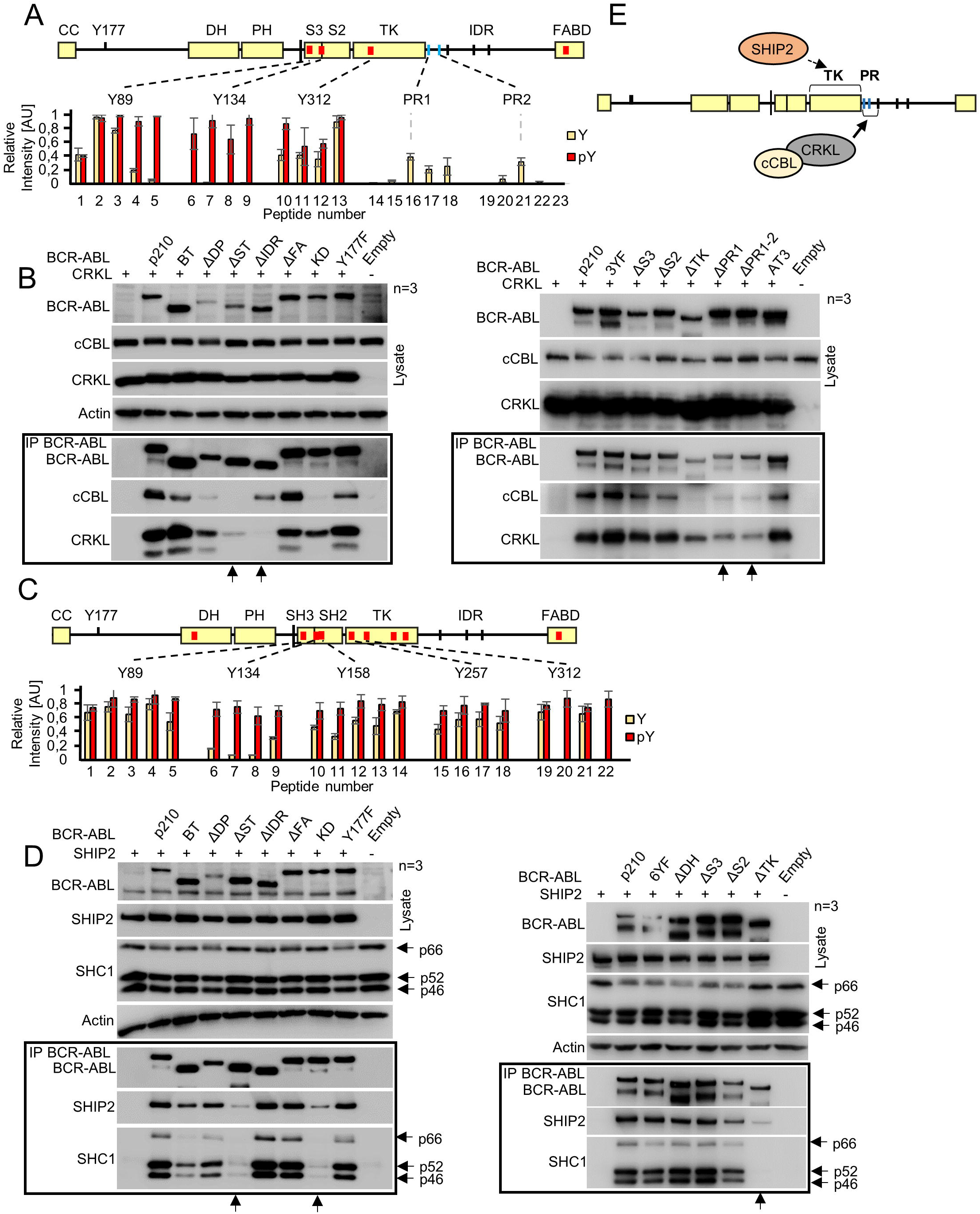
Interaction of CRKL and SHIP2 with BCR-ABL. (A) BCR-ABL scheme with red lines indicating potential binding sites identified by microarray. Graph shows binding intensities for phosphorylated (red) and non-phosphorylated peptides; strong binding is shown for phosphorylated Y89, Y134 and Y312. (B) Immunoprecipitation of BCR-ABL with CRKL in transfected 293T cells. Deletion of IDR and both PR1 and PR2 sites (constructs ΔPR1 and ΔPR1-2) largely limits CRKL co-IP with BCR-ABL (arrows). Substituting Y89, Y134 and Y312 to phenylalanines (3YF) produced no effect on CRKL interaction with BCR-ABL. cCBL also binds less to mutants lacking PR1 and PR2 sites, suggesting at least some proportion of cCBL binds to BCR-ABL via CRKL. (C) Microarray analysis of SHIP2 binding to BCR-ABL shows multiple binding sites in the SH3-SH2-TK domains, association with the tyrosine-phosphorylated motifs is indicated in red. (D) Immunoprecipitation of BCR-ABL constructs with SHIP2 in transfected 293T cells. Deletion of SH3-SH2-TK’(ΔST), or TK domain (ΔTK) largely limits SHIP2 binding (arrows). Substituting Y89, Y134, Y158, Y257, Y312 and Y469 to phenylalanines (Y6F) had no effect on SHIP2 binding. SHC1 associated with BCR-ABL in a manner similar to SHIP2, suggesting mutual interaction. (E) Scheme of proposed interaction. CRKL binds to region containing PR1 and PR2 and partially mediates cCBL binding. SHIP2 binds to the TK region of ABL. Data are representative for three independent experiments (n=3).

**Figure 7.**
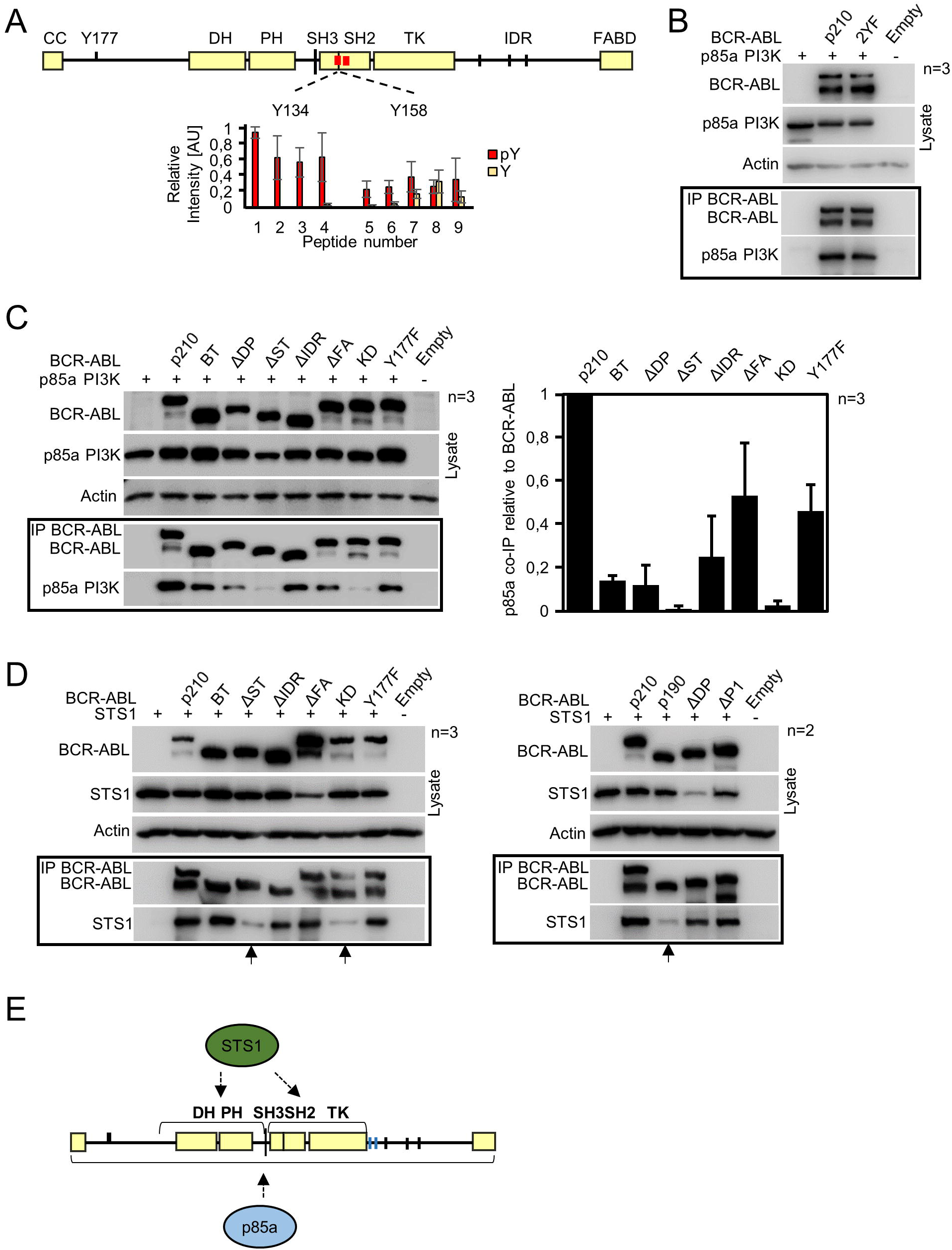
Analysis of p85a-PI3K and STS1 interaction with BCR-ABL. (A) BCR-ABL scheme with potential p85a-PI3K binding sites identified by peptide microarray. Graph shows intensities for phosphorylated (red) and non-phosphorylated peptides containing Y134 and Y158. (B) Immunoprecipitation (IP) of BCR-ABL with p85a-PI3K in 293T cells. Deletion of SH3, SH2 and TK domains largely limits p85a-PI3K interaction with BCR-ABL (arrows). Substitutions of Y134 and Y158 to phenylalanines (2YF) had no effect on p85a-PI3K interaction with BCR-ABL. (C) Immunoprecipitation of BCR-ABL with STS1. Deletion of SH3, SH2, and TK domains largely abrogates BCR-ABL effect on interaction with STS1. Deletion of DH-PH domains in p190 and ΔDP also limits binding of STS1. (D) Scheme of the proposed interaction. p85a-PI3K interacts with SH3/ SH2/TK domain of BCR-ABL, whereas STS1 requires also the PH domain for association. Data are representative for three independent experiments (n=3).

SHIP2 is an inositol 5’phosphatase whose catalytic activity removes 5’phosphate from phosphatidyl inositol triphosphates PI(3,4,5)P_3_, converting them into PI(3,4)P_2_. SHIP2 has role in signaling after activation of hematopoietic growth factor receptors leading to its phosphorylation and SHC1 association (Wisniewski et al. 1999; Srivastava, Sudan, and Kerr 2013). Depending on the cellular context, SHIP2 can either be anti or pro-oncogenic (Taylor et al. 2000; Hoekstra et al. 2016), however its role in CML has not been established. SHIP2 is constitutively phosphorylated in both primary CML cells and p210 expressing cells (Odai et al. 1997; Wisniewski et al. 1999). Phosphorylated SHIP2 was found to bind SH3 domain of BCR-ABL but not the SH2 domain (Wisniewski et al. 1999). Our microarray data showed multitude of potential binding sites mostly falling into ordered domains of BCR-ABL, including SH3-SH2-TK domains, and phosphorylated tyrosines, Y89, Y134, Y158, Y257, Y312 and Y469 located within these domains (Fig. 6C). To test these putative binding sites, we co-expressed BCR-ABL constructs with SHIP2 and verified if SHIP2 co-immunoprecipitated with BCR-ABL. Deletion of SH3-SH2-TK domains partially abolished interaction with SHIP2, similar the BCR-ABL-KD mutant (Fig. 6D; arrows). The Y177F substitution in BCR-ABL, as well as deletion of DH-PH, IDR and FABD domains produced no effect on interaction with SHIP2. Individual deletions of SH3 and SH2 (ΔS3, ΔS2) had no effect on SHIP2 association with BCR-ABL, in contrast to deletion of TK domain, which abolished the binding almost completely (Fig. 6D; arrows). Finally, to verify if the interaction is mediated by phosphorylated tyrosines within SH3-SH2-TK domains, we created mutant BCR-ABL with 6 tyrosines selected by microarray were mutated to phenylalanines (Y89F, Y134F, Y158F, Y257F, Y312F and Y469F; BCR-ABL-6YF). SHIP2 and BCR-ABL-6YF co-immunoprecipitated normally. Because SHIP2 is known to bind SHC1 (Wisniewski et al. 1999), we evaluated whether binding of SHIP2 to BCR-ABL is dependent on SHC binding. Their binding to BCR-ABL deletion constructs shows similar pattern, suggesting that they may bind cooperatively. However, ΔTK construct can still partially bind SHIP2, but no SHC1, suggesting that SHIP2 binding is not entirely dependent on SHC1 (Fig. 6D).

### Interaction of p85a-PI3K and STS1 with BCR-ABL

p85a is a small regulatory subunit of phosphatidyl inositol 3-kinase (PI3K). Inhibiting p85a-PI3K expression or PI3K activity leads to inhibition of growth in BCR-ABL positive cells, identifying p85a-PI3K as important interactor (Skorski et al. 1995, 1997). p85a-PI3K contains SH3, RhoGAP and two C-terminal SH2 domains. Consensus binding motif for p85a-PI3K is phosphorylated YxxM motif (S. Zhou et al. 1993) and its binding by SH2 domain of p85a-PI3K leads to activation of PI3K activity in cells (Escobedo et al. 1991; Holt et al. 1994). Notably, there is one YxxM motif present in BCR-ABL TK domain involving Tyr469, however its mutation showed no effect on BCR-ABL association with p85a-PI3K or PI3K activation (Jain et al. 1996). Our microarray analysis revealed two potential binding sites that were only five residues apart and included phosphotyrosines Y134 in the SH3 and Y158 in SH2 domain (Fig. 7A). To verify if these two tyrosines mediate binding of p85a-PI3K to BCR-ABL, we created double mutant with Y134 and Y158 mutated to phenylalanines (2YF). This mutant had no effect on p85a-PI3K binding (Fig. 7B), however, co-immunoprecipitations with basic deletion toolkit revealed decrease in p85a binding to virtually all constructs. Binding was almost abolished by ΔST and KD mutant. Deletion of N-terminal part also showed decrease in p85a binding, again likely to be caused by lack of kinase activity. Y177F substitution showed about 50% decrease in p85 binding, Deletion of DH and PH domains abrogated interaction with p85a by cca 80%, deletion of IDR by 70% and deletion of FABD domain by 40% (Fig. 7C). This complex mode of binding to BCR-ABL may reflect the fact that p85 has ability to bind multiple components within the BCR-ABL complex, such as CBL, SHC1 and GAB2 (S. Ren et al. 2005).

STS1 is a tyrosine phosphatase that was recently found to facilitate BCR-ABL dephosphorylation and thus appears to be negative regulator of BCR-ABL kinase activity (Mian et al. 2019). It has UBA domain, SH3 and protein tyrosine phosphatase domain. Proteomic screens revealed that STS1 has high preference for p210 BCR-ABL over p190 BCR-ABL, that lacks DH and PH domains (Reckel et al. 2017; Cutler et al. 2017). We did not detect any signals indicative of interaction between STS1 and BCR-ABL on microarrays, despite optimization and use of two different recombinant STS1 (not shown). Co-immunoprecipitation experiments show that kinase activity of BCR-ABL is essential for STS1 binding, as BCR-ABL-KD and deletion of SH3-SH2-TK domains completely abolished STS1 binding (Fig. 7D). Interaction of STS1 with p190 BCR-ABL lacking DH and PH domains is also largely limited. However, smaller deletions of these domains ΔDP and deletion of PH domain ΔP1 do not limit interaction with STS1, suggesting that the entire region missing in p190 (aa 425-927) is necessary for STS1 interaction.

### SHIP2 is required for BCR-ABL-mediated phosphorylation of SHC1

In order to elucidate the role of SHIP2 in BCR-ABL mediated signaling, we abolished endogenous SHIP2 expression by CRISPR/Cas9 in 293T cells (Ship2^Crispr^ cells), transfected these cells with BCR-ABL, and determined the effect of SHIP2 loss on BCR-ABL-mediated signaling. In contrast to wildtype 293T cells, Ship2^Crispr^ cells could not phosphorylate SHC1 on tyrosines 239 and 240 (Fig. 8A). Addition of wildtype SHIP2 back into Ship2^Crispr^ cells rescued SHC1 phosphorylation. However, co-transfecting SHIP2 with inactivated phosphatase activity (PD) only partially rescued SHC1 phosphorylation, suggesting that SHIP2 catalytic activity is important for BCR-ABL-mediated SHC1 phosphorylation (Fig. 8B). As SHIP2 and SHC1 are known interactors (Wisniewski et al. 1999), we asked if SHC1 will interact with BCR-ABL in Ship2^Crispr^ cells. SHC1 interacted with BCR-ABL normally in Ship2^Crispr^ cells (Fig. 8C).

**Figure 8.**
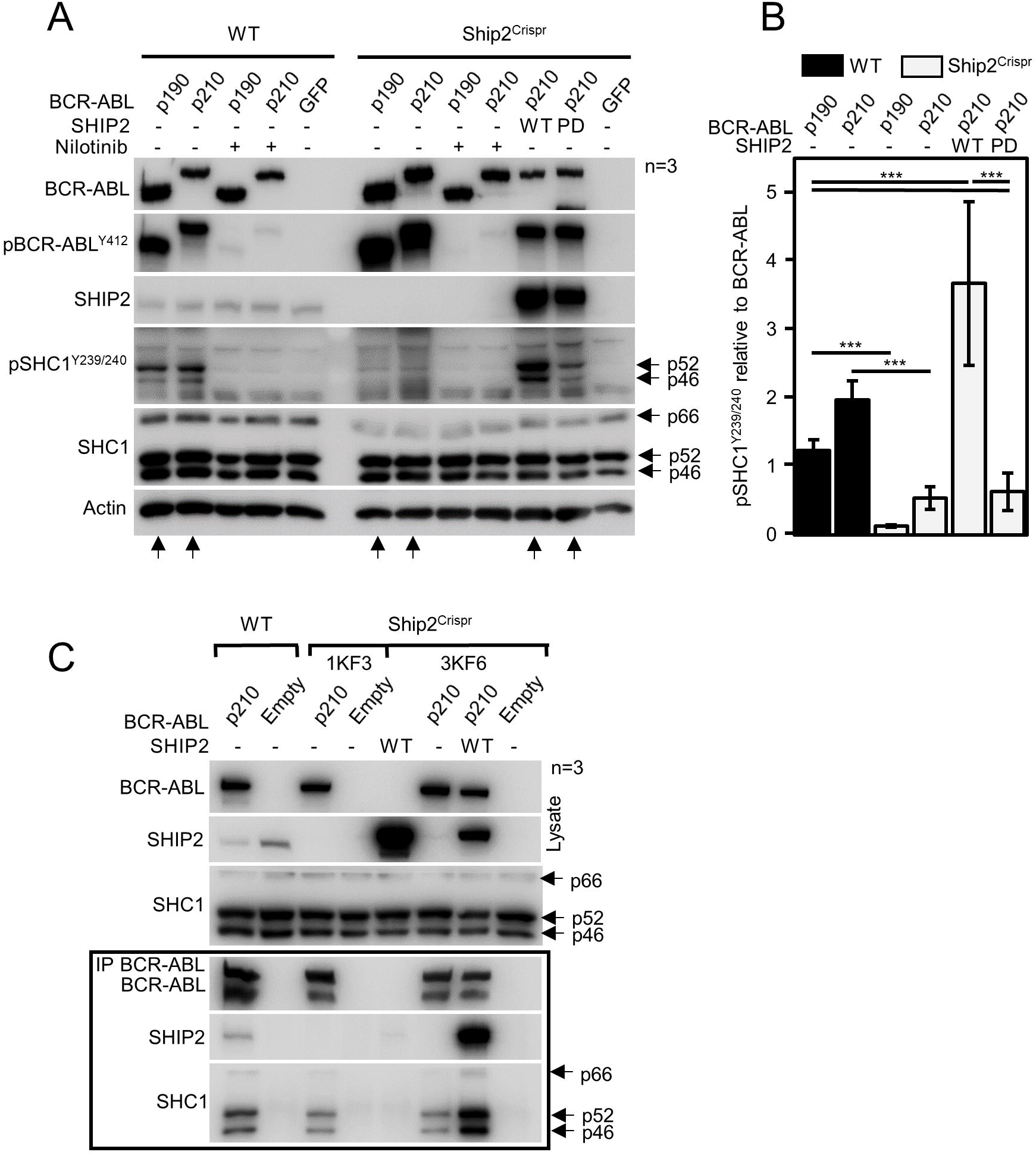
SHIP2 is required for BCR-ABL-mediated phosphorylation of SHC1. (A) Wildtype (wt) 293T cells and cells with SHIP2 deleted by CRISPR/Cas9 (Ship2^Crispr^) 293T were transfected by BCR-ABL, and analyzed for presence and phosphorylation (p) of given proteins by westernblot. In contrast to wildtype cells, Ship2^Crispr^ cells could not phosphorylate SHC1. Addition of wt SHIP2 into the Ship2^Crispr^ 293T cells rescued BCR-ABL-mediated SHC1 phosphorylation, while addition of catalytically inactive (phosphatase-dead, PD) had no effect. Same results were obtained with two independent Ship2^Crispr^ cell lines. (B) Quantification of SHC1 phosphorylation normalized to BCR-ABL levels. Bars represent averages from 8 measurements with indicated S.D., statistically significant differences are indicated (Student’s *t*-test, \*\*\**p*<0.001). (C) SHC1 interacts with BCR-ABL in Ship2^Crispr^ 293T cells. Wt and Ship2^Crispr^ 293T cells (clones 1KF3 and 3KF6) were transfected with BCR-ABL followed by BCR-ABL immunoprecipitation (IP). SHC1 co-immunoprecipitates with BCR-ABL in Ship2^Crispr^, suggesting that lack of SHC1 phosphorylation in Ship2^Crispr^ cells is not due to loss of interaction with BCR-ABL. Data are representative for three independent experiments (n=3).

We present here detailed characterization of BCR-ABL complex. All of the proteins evaluated were sensitive to kinase activity of BCR-ABL and with the exception of CRKL, their binding to KD BCR-ABL was severely reduced. It is likely that at least some interactors need to be phosphorylated in order to bind BCR-ABL, as processive phosphorylation of substrate followed by its binding to kinase is known phenomenon. Alternatively, BCR-ABL can phosphorylate its own residues that can pose binding sites for the interactors.

Our data show that GRB2 directly binds to phosphorylated Y177, which is necessary and sufficient for binding, as Y177F substitution abolishes binding. SOS1 also does not interact with Y177F BCR-ABL, however it is known interactor of GRB2 and therefore we did not analyze its binding by microarrays. Interestingly, cCBL also largely depends on Y177 for its binding to BCR-ABL. cCBL is also known CRKL interactor and deletion of CRKL binding sites PR1 and PR2 within IDR region also largely limited binding of cCBL. Lack of interaction of GRB2, SOS1 and cCBL with KD BCR-ABL and ΔTK can likely be attributed to lack of autophosphorylation of Y177 and/or the interactors.

For the remaining interactors we did not find as clearly defined binding site as for GRB2 and SOS1, as entire domains or larger epitopes of BCR-ABL were necessary to delete to limit binding. Deletion of DH and PH domains limited binding of STS1 and p85a PI3K and deletion of PH domain limited SHC1 binding. We found that CRKL requires proline rich regions in IDR for binding and that SHIP2 requires TK domain. Furthermore, we uncovered that SHIP2 mediates SHC1 phosphorylation in the BCR-ABL complex and that SHIP2 phosphatase activity is largely responsible for the mediation of the phosphorylation. Therefore, SHIP2 phosphatase activity may paradoxically contribute to Ras activation by SHC1.

## Material and methods

### Cell culture, vectors, transfection and CRISPR/Cas9

293T cells and NIH3T3 cells were obtained from ATCC, and propagated in DMEM media, supplemented with 10% FBS and antibiotics (ThermoFisher). All expression vectors are listed in Table S1. Cells were transiently transfected using FuGENE HD, according to the manufacturer’s protocol (Promega). p210 isoform b3a2 and p190 BCR-ABL sequences (isoform b3a2) were cloned into pCR3.1 vector with N-terminal FLAG tag. Polymerase chain reaction (PCR) mutagenesis was used to generate all BCR-ABL variants. GRB2 (RC200469), p85a-PI3K (RG210544) and SHIP2 (RC214716) vectors were obtained from Origene, CrkL (HG11261-CH) and Sts1 (HG13868-NF) vectors were obtained from Sino Biological. Their coding sequences were subcloned into modified pCMV6 entry vector, where C-terminal Myc-DDK tag was replaced by HIS-V5 tag using a NEBuilder HiFi DNA assembly kit (New England Biolabs). SHIP2 deletion in 293T cells was carried out by CRISPR/Cas9 technology ***(Ran et al. 2013)***. CHOPCHOP tool was used to design sgRNAs for a pair of SpCas9n (D10A) nickases, which targeted 5’-CGATGGCAGCTTCCTGGTCC-3’, 5’- GCGCTCTGCGTCCTGTGAGT-3’ sites in the first exon of the SHIP2 gene ***(Montague et al. 2014)***. Successful targeting disrupted open reading frame of SHIP2, which was detected as loss of SHIP2 in individual clones by western blot. Targeted locus was PCR amplified using 5’-CCGGGCGGCCGCGGAGGAG-3’, 5’-TCTGGCGTCCCACCGCCCCAGAAAC-3’, inserted into pGEM T-Easy vector (Promega) and sequenced for determination of SHIP2 genotype.

### Western blot and immunoprecipitation (IP)

About 5×10^6^ of transfected cells were lysed for 30 minutes at 4°C in 1 ml of IP lysis buffer containing 50 mM tris-HCl (pH 7.4), 150 mM NaCl, 0.5% NP-40, 2 mM EDTA, 1 mM Na_3_VO_4_ and protease inhibitors (Roche). For IP, 25 ul of Dynabeads protein G (ThermoFisher) were bound to FLAG antibody (F1804, Sigma-Aldrich) or V5 antibody (R96025, Invitrogen) according to the manufacturer’s protocol. Immunocomplexes were collected overnight at 4°C. Proteins attached to the beads were eluted to 60 ul of 2xLaemmli buffer. For western blot, cell lysates or immunoprecipitates were resolved by SDS-PAGE, transferred onto a PVDF membrane, and visualized by chemiluminiscence using Pierce ECL (Thermo Fisher Scientific), Immobilon Western (Millipore), Clarity (Biorad) or SuperSignal West Femto (Thermo Fisher Scientific) substrates. Table S2 lists all antibodies used in the study.

### Gradient ultracentrifugation, BN-PAGE and proximity ligation assay (PLA)

The gradient ultracentrifugation was done as described before (Kunova Bosakova et al. 2019). Briefly, the native lysates (50 mM Tris-HCl pH 7.4, 150 mM NaCl, 0.5 % Igepal CA-630, 1 mM EDTA pH 8, 0.25 % sodium deoxycholate, 1 mM Na_2_VO_4_; proteinase inhibitors) were cleared, loaded on 15-40% sucrose gradient (1 mM Tris-HCl pH 7.4, 150 mM NaCl, 3 mM MgCl_2_; proteinase inhibitors), and centrifuged at 40,000 rpm/4 °C/16 hours using SW 40 Ti rotor (Beckman Coulter). Approximately 18 fractions were collected from each gradient, the proteins were precipitated with 10% TCA, dissolved in 2xLaemmli buffer and boiled. The respective fractions, as well as the cell lysates collected before ultracentrifugation were resolved by western blot, and co-sedimentation of the endogenous interaction partners with expressed BCR-ABL was analyzed by densitometry (ImageJ; http://imagej.nih.gov/ij/). To quantify western blots in shown in Fig. 1A, the relative abundance of each of the proteins in the dominant p190 BCR-ABL fractions (#6 in control and #5 in nilotinib treated cells) was calculated by densitometry of all fractions, and plotted. As only proteins saturated on BCR-ABL were analyzed, any peaks exceeding the relative optical density of 1 (red dashed lines) after treatment with nilotinib would suggest dissociation of the protein from the BCR-ABL complex, as observed with GRB2. Analogically, the peak of p85a-PI3K above 1 in control cells and absence of the protein in heavier BCR-ABL fractions suggest that not all BCR-ABL complexes involve p85a-PI3K. The BN-PAGE was carried out as described before (Kunova Bosakova et al. 2019). The native cell lysates have been loaded on 4-15% native gels, after the native electrophoresis the lane sample strips were excised from the gel, denatured and resolved in 12% SDS-PAGE gels. For Duolink® PLA (Sigma), cells were fixed in paraformaldehyde, post-fixed in ice-cold methanol and stained according to manufacturer’s protocol. Mouse FLAG (F1804; Sigma) and Goat V5 (sc-83849; Santa Cruz) antibodies were used for PLA; rabbit anti-c-ABL (2862S; Cell Signaling) was used to counterstain the transfected cells. Secondary antibodies conjugated with AlexaFluor488/594 were from Invitrogen. PLA counting analysis was done in Fiji (http://fiji.sc/Fiji) using maximum projections of Z-stacks.

### Peptide microarrays

Pepstar™ microarray technology (JPT Peptide Technologies GmbH, Germany) was used to identify sites in BCR-ABL involved in binding of individual core complex interactors. Peptide library was generated by dividing entire p210 BCR-ABL protein sequence into 13aa long peptides (674) with 10 residue overlap between neighbouring peptides. Peptides were synthesized and immobilized on a glass slide. To account for phosphorylation at known BCR-ABL sites, namely Y177, Y328, Y360 in BCR and Y89, Y134, Y147, Y158, Y191, Y204, Y234, Y251, Y272, Y276, Y312, Y412, S637-638, T735 in ABL, phosphorylated versions of the 94 corresponding peptides were also included in the microarrays. For each experiment, experimental and control microarrays were processed in parallel. Arrays were incubated with recombinant interactors. GRB2 (TP300469), SHC1 (TP304362), STS1 (TP303523), CRKL (TP308129) and cCbl (TP314069) were obtained from Origene, SHIP2 (P09-20G-10) and p85a-PI3K (P31-30H) were obtained from SignalChem. Signal was developed by incubation of arrays with primary antibodies against GST (G1160), FLAG (F1804, Sigma-Aldrich) and HIS (sc-8036, Santa-Cruz) and secondary, Cy-5-coupled antibody (715-175-151, Jackson ImmunoResearch). Arrays were scanned using InnoScan 1100 AL fluorescence scanner and data were analyzed by Mapix software (Innopsys). Each microarray slide contained three entire peptide libraries, which were analyzed as technical replicates. Fluorescence intensities from were plotted as a function of the peptide number (BCR-ABL primary sequence). First, the intensities from all spots in the control microarray were used to calculate arithmetic average (∅_c_) and the standard deviation (σ_c_). If a signal intensity for a peptide spot in the control microarray exceeded value of (∅_c_+1.σ_c_) in all three replicates, the respective peptide (microarray spot) was excluded from further analysis. Subsequently, fluorescence intensities in negative control microarrays were subtracted from values in experimental microarrays and these values were used for all further analyses. To allow direct comparison of fluorescence intensities from the three technical replicates, relative fluorescence intensities for each peptide were calculated by dividing each signal by biggest value in respective microarray. Finally, relative fluorescence intensities from the three replicates were averaged, plotted as a function of the peptide number (BCR-ABL primary sequence) and arithmetic average (∅_ER_) and standard deviation (σ_ER_) were calculated from the averaged fluorescence intensities. Potential binding site in the microarray was considered when at least three consecutive peptides exceeded (∅_ER_+1.σ_ER_) in all replicates.

## Acknowledgement

This study was supported by Agency for Healthcare Research of the Czech Republic, project no. 15-34405A (CEP: NV15-34405A).

## References

Bewry, N. N., R. R. Nair, M. F. Emmons, D. Boulware, J. Pinilla-Ibarz, and L. A. Hazlehurst. 2008. “Stat3 Contributes to Resistance toward BCR-ABL Inhibitors in a Bone Marrow Microenvironment Model of Drug Resistance.” Molecular Cancer Therapeutics 7 (10): 3169–75. https://doi.org/10.1158/1535-7163.MCT-08-0314.

Brasher, B. B., S. Roumiantsev, and R. A. Van Etten. 2001. “Mutational Analysis of the Regulatory Function of the C-Abl Src Homology 3 Domain.” Oncogene 20 (53): 7744–52. https://doi.org/10.1038/sj.onc.1204978.

Brehme, Marc, Oliver Hantschel, Jacques Colinge, Ines Kaupe, Melanie Planyavsky, Thomas Köcher, Karl Mechtler, Keiryn L. Bennett, and Giulio Superti-Furga. 2009. “Charting the Molecular Network of the Drug Target Bcr-Abl.” Proceedings of the National Academy of Sciences 106 (18): 7414–7419.

Buday, László, Asim Khwaja, Szabolcs Sipeki, Anna Faragó, and Julian Downward. 1996. “Interactions of Cbl with Two Adaptor Proteins, Grb2 and Crk, upon T Cell Activation.” Journal of Biological Chemistry 271 (11): 6159–6163.

Cilloni, Daniela, and Giuseppe Saglio. 2012. “Molecular Pathways: BCR-ABL.” Clinical Cancer Research: An Official Journal of the American Association for Cancer Research 18 (4): 930–37. https://doi.org/10.1158/1078-0432.CCR-10-1613.

Corbin, Amie S., Anupriya Agarwal, Marc Loriaux, Jorge Cortes, Michael W. Deininger, and Brian J. Druker. 2011. “Human Chronic Myeloid Leukemia Stem Cells Are Insensitive to Imatinib despite Inhibition of BCR-ABL Activity.” Journal of Clinical Investigation 121 (1): 396–409. https://doi.org/10.1172/JCI35721.

Cortes, J.E., D.-W. Kim, J. Pinilla-Ibarz, P. le Coutre, R. Paquette, C. Chuah, F.E. Nicolini, et al. 2013. “A Phase 2 Trial of Ponatinib in Philadelphia Chromosome–Positive Leukemias.” New England Journal of Medicine 369 (19): 1783–96. https://doi.org/10.1056/NEJMoa1306494.

Cutler, J A, R Tahir, S K Sreenivasamurthy, C Mitchell, S Renuse, R S Nirujogi, A H Patil, et al. 2017. “Differential Signaling through P190 and P210 BCR-ABL Fusion Proteins Revealed by Interactome and Phosphoproteome Analysis.” Leukemia 31 (7): 1513–24. https://doi.org/10.1038/leu.2017.61.

Donovan, J. A., R. L. Wange, W. Y. Langdon, and L. E. Samelson. 1994. “The Protein Product of the C-Cbl Protooncogene Is the 120-KDa Tyrosine-Phosphorylated Protein in Jurkat Cells Activated via the T Cell Antigen Receptor.” Journal of Biological Chemistry 269 (37): 22921–24.

Druker, B. J., S. Tamura, E. Buchdunger, S. Ohno, G. M. Segal, S. Fanning, J. Zimmermann, and N. B. Lydon. 1996. “Effects of a Selective Inhibitor of the Abl Tyrosine Kinase on the Growth of Bcr-Abl Positive Cells.” Nature Medicine 2 (5): 561–66.

Eide, Christopher A., Lauren T. Adrian, Jeffrey W. Tyner, Mary Mac Partlin, David J. Anderson, Scott C. Wise, Bryan D. Smith, et al. 2011. “The ABL Switch Control Inhibitor DCC-2036 Is Active against the Chronic Myeloid Leukemia Mutant BCR-ABLT315I and Exhibits a Narrow Resistance Profile.” Cancer Research 71 (9): 3189–95. https://doi.org/10.1158/0008-5472.CAN-10-3224.

Escobedo, J. A., D. R. Kaplan, W. M. Kavanaugh, C. W. Turck, and L. T. Williams. 1991. “A Phosphatidylinositol-3 Kinase Binds to Platelet-Derived Growth Factor Receptors through a Specific Receptor Sequence Containing Phosphotyrosine.” Molecular and Cellular Biology 11 (2): 1125–1132.

Goga, Andrei, Jami McLaughlin, Daniel EH Afar, Douglas C. Saffran, and Owen N. Witte. 1995. “Alternative Signals to RAS for Hematopoietic Transformation by the BCR-ABL Oncogene.” Cell 82 (6): 981–988.

Gorre, M. E., M. Mohammed, K. Ellwood, N. Hsu, R. Paquette, P. N. Rao, and C. L. Sawyers. 2001. “Clinical Resistance to STI-571 Cancer Therapy Caused by BCR-ABL Gene Mutation or Amplification.” Science (New York, N.Y.) 293 (5531): 876–80. https://doi.org/10.1126/science.1062538.

Grebien, Florian, Oliver Hantschel, John Wojcik, Ines Kaupe, Boris Kovacic, Arkadiusz M. Wyrzucki, Gerald D. Gish, et al. 2011. “Targeting the SH2-Kinase Interface in Bcr-Abl Inhibits Leukemogenesis.” Cell 147 (2): 306–19. https://doi.org/10.1016/j.cell.2011.08.046.

Hazlehurst, Lori A., Nadine N. Bewry, Rajesh R. Nair, and Javier Pinilla-Ibarz. 2009. “Signaling Networks Associated with BCR-ABL-Dependent Transformation.” Cancer Control: Journal of the Moffitt Cancer Center 16 (2): 100–107. https://doi.org/10.1177/107327480901600202.

Heaney, Conor, Kathryn Kolibaba, Arun Bhat, Tsukasa Oda, Sayuri Ohno, Shane Fanning, and Brian J. Druker. 1997. “Direct Binding of CRKL to BCR-ABL Is Not Required for BCR-ABL Transformation.” Blood 89 (1): 297–306.

Hochhaus, A., M. Baccarani, M. Deininger, J. F. Apperley, J. H. Lipton, S. L. Goldberg, S. Corm, et al. 2008. “Dasatinib Induces Durable Cytogenetic Responses in Patients with Chronic Myelogenous Leukemia in Chronic Phase with Resistance or Intolerance to Imatinib.” Leukemia 22 (6): 1200–1206. https://doi.org/10.1038/leu.2008.84.

Hoekstra, Elmer, Asha M. Das, Marcella Willemsen, Marloes Swets, Peter JK Kuppen, Christien J. van der Woude, Marco J. Bruno, Jigisha P. Shah, Timo LM ten Hagen, and John D. Chisholm. 2016. “Lipid Phosphatase SHIP2 Functions as Oncogene in Colorectal Cancer by Regulating PKB Activation.” Oncotarget 7 (45): 73525.

Hoeve, Johanna ten, Ralph B. Arlinghaus, Jie Qiang Guo, Nora Heisterkamp, and John Groffen. 1994. “Tyrosine Phosphorylation of CRKL in Philadelphia+ Leukemia.” Blood 84 (6): 1731–1736.

Holt, Kathleen H., L. Olson, W. Scott Moye-Rowley, and Jeffrey E. Pessin. 1994. “Phosphatidylinositol 3-Kinase Activation Is Mediated by High-Affinity Interactions between Distinct Domains within the P110 and P85 Subunits.” Molecular and Cellular Biology 14 (1): 42–49.

Jain, Suresh K., M. Susa, Marilyn L. Keeler, Nadia Carlesso, Brian Druker, and Lyuba Varticovski. 1996. “PI 3-Kinase Activation in BCR/Abl-Transformed Hematopoietic Cells Does Not Require Interaction of P85 SH2 Domains with P210 BCR/Abl.” Blood 88 (5): 1542–1550.

Kunova Bosakova, Michaela Alexandru Nita, Tomas Gregor, Miroslav Varecha, Iva Gudernova, Bohumil Fafilek, Tomas Barta, et al. 2019. “Fibroblast Growth Factor Receptor Influences Primary Cilium Length through an Interaction with Intestinal Cell Kinase.” Proceedings of the National Academy of Sciences 116 (10): 4316–25. https://doi.org/10.1073/pnas.1800338116.

Lewitzky, M., C. Kardinal, N. H. Gehring, E. K. Schmidt, B. Konkol, M. Eulitz, W. Birchmeier, U. Schaeper, and S. M. Feller. 2001. “The C-Terminal SH3 Domain of the Adapter Protein Grb2 Binds with High Affinity to Sequences in Gab1 and SLP-76 Which Lack the SH3-Typical P-x-x-P Core Motif.” Oncogene 20 (9): 1052–62. https://doi.org/10.1038/sj.onc.1204202.

Meisner, Herman, Bruce R. Conway, David Hartley, and Michael P. Czech. 1995. “Interactions of Cbl with Grb2 and Phosphatidylinositol 3’-Kinase in Activated Jurkat Cells.” Molecular and Cellular Biology 15 (7): 3571–3578.

Meyn, Malcolm A., Matthew B. Wilson, Fadi A. Abdi, Nathalie Fahey, Anthony P. Schiavone, Jiong Wu, James M. Hochrein, John R. Engen, and Thomas E. Smithgall. 2006. “Src Family Kinases Phosphorylate the Bcr-Abl SH3-SH2 Region and Modulate Bcr-Abl Transforming Activity.” Journal of Biological Chemistry 281 (41): 30907–16. https://doi.org/10.1074/jbc.M605902200.

Mian, Afsar A., Ines Baumann, Marcus Liebermann, Florian Grebien, Giulio Superti-Furga, Martin Ruthardt, Oliver G. Ottmann, and Oliver Hantschel. 2019. “The Phosphatase UBASH3B/Sts-1 Is a Negative Regulator of Bcr-Abl Kinase Activity and Leukemogenesis.” Leukemia, April, 1. https://doi.org/10.1038/s41375-019-0468-y.

Mitra, Aninda, Kotagiri Sasikumar, B.V.V. Parthasaradhi, and Vegesna Radha. 2013. “The Tyrosine Phosphatase TC48 Interacts with and Inactivates the Oncogenic Fusion Protein BCR-Abl but Not Cellular Abl.” Biochimica et Biophysica Acta (BBA) - Molecular Basis of Disease 1832 (1): 275–84. https://doi.org/10.1016/j.bbadis.2012.10.014.

Modugno, Michele. 2014. “New Resistance Mechanisms for Small Molecule Kinase Inhibitors of Abl Kinase.” Drug Discovery Today: Technologies 11 (March): 5–10. https://doi.org/10.1016/j.ddtec.2013.12.001.

Montague, Tessa G., José M. Cruz, James A. Gagnon, George M. Church, and Eivind Valen. 2014. “CHOPCHOP: A CRISPR/Cas9 and TALEN Web Tool for Genome Editing.” Nucleic Acids Research 42 (Web Server issue): W401–407. https://doi.org/10.1093/nar/gku410.

Nichols, Gwen L., M. A. Raines, J. Carlos Vera, L. Lacomis, P. a Tempst, and D. W. Golde. 1994. “Identification of CRKL as the Constitutively Phosphorylated 39-KD Tyrosine Phosphoprotein in Chronic Myelogenous Leukemia Cells.” Blood 84 (9): 2912–2918.

Oda, Tsukasa, Conor Heaney, John R. Hagopian, Keiko Okuda, James D. Griffin, and Brian J. Druker. 1994. “Crkl Is the Major Tyrosine-Phosphorylated Protein in Neutrophils from Patients with Chronic Myelogenous Leukemia.” Journal of Biological Chemistry 269 (37): 22925–22928.

Odai, Hideharu, Ko Sasaki, Akihiro Iwamatsu, Tetsuya Nakamoto, Hiroo Ueno, Tetsuya Yamagata, Kinuko Mitani, Yoshio Yazaki, and Hisamaru Hirai. 1997. “Purification and Molecular Cloning of SH2-and SH3-Containing Inositol Polyphosphate-5-Phosphatase, Which Is Involved in the Signaling Pathway of Granulocyte-Macrophage Colony-Stimulating Factor, Erythropoietin, and Bcr-Abl.” Blood 89 (8): 2745–2756.

Panchamoorthy, Govindaswamy, Toru Fukazawa, Sachiko Miyake, Stephen Soltoff, Kris Reedquist, Brian Druker, Steve Shoelson, Lewis Cantley, and Hamid Band. 1996. “P120 Is a Major Substrate of Tyrosine Phosphorylation upon B Cell Antigen Receptor Stimulation and Interacts in Vivo with Fyn and Syk Tyrosine Kinases, Grb2 and Shc Adaptors, and the P85 Subunit of Phosphatidylinositol 3-Kinase.” Journal of Biological Chemistry 271 (6): 3187–3194.

Pendergast, Ann Marie, Lawrence A. Quilliam, Larry D. Cripe, Craig H. Bassing, Zonghan Dai, Nanxin Li, Andreas Batzer, et al. 1993. “BCR-ABL-Induced Oncogenesis Is Mediated by Direct Interaction with the SH2 Domain of the GRB-2 Adaptor Protein.” Cell 75 (1): 175–185.

Preyer, Martin, Paolo Vigneri, and Jean Y. J. Wang. 2011. “Interplay between Kinase Domain Autophosphorylation and F-Actin Binding Domain in Regulating Imatinib Sensitivity and Nuclear Import of BCR-ABL.” Edited by Gen Sheng Wu. PLoS ONE 6 (2): e17020. https://doi.org/10.1371/journal.pone.0017020.

Ran, F. Ann, Patrick D. Hsu, Jason Wright, Vineeta Agarwala, David A. Scott, and Feng Zhang. 2013. “Genome Engineering Using the CRISPR-Cas9 System.” Nature Protocols 8 (11): 2281–2308. https://doi.org/10.1038/nprot.2013.143.

Reckel, S, R Hamelin, S Georgeon, F Armand, Q Jolliet, D Chiappe, M Moniatte, and O Hantschel. 2017. “Differential Signaling Networks of Bcr–Abl P210 and P190 Kinases in Leukemia Cells Defined by Functional Proteomics.” Leukemia 31 (7): 1502–12. https://doi.org/10.1038/leu.2017.36.

Ren, Ruibao, Zheng-Sheng Ye, and David Baltimore. 1994. “Abl Protein-Tyrosine Kinase Selects the Crk Adapter as a Substrate Using SH3-Binding Sites.” Genes & Development 8 (7): 783–795.

Ren, Shu-yue, Fan Xue, Jan Feng, and Tomasz Skorski. 2005. “Intrinsic Regulation of the Interactions between the SH3 Domain of P85 Subunit of Phosphatidylinositol-3 Kinase and the Protein Network of BCR/ABL Oncogenic Tyrosine Kinase.” Experimental Hematology 33 (10): 1222–28. https://doi.org/10.1016/j.exphem.2005.06.030.

Rousselot, Philippe, Aude Charbonnier, Pascale Cony-Makhoul, Philippe Agape, Franck E. Nicolini, Bruno Varet, Martine Gardembas, et al. 2014. “Loss of Major Molecular Response as a Trigger for Restarting Tyrosine Kinase Inhibitor Therapy in Patients with Chronic-Phase Chronic Myelogenous Leukemia Who Have Stopped Imatinib after Durable Undetectable Disease.” Journal of Clinical Oncology: Official Journal of the American Society of Clinical Oncology 32 (5): 424–30. https://doi.org/10.1200/JCO.2012.48.5797.

Sattler, Martin, M. Golam Mohi, Yuri B. Pride, Laura R. Quinnan, Nicole A. Malouf, Klaus Podar, Franck Gesbert, et al. 2002. “Critical Role for Gab2 in Transformation by BCR/ABL.” Cancer Cell 1 (5): 479–492.

Senechal, Kristen, Jocelyn Halpern, and Charles L. Sawyers. 1996. “The CRKL Adaptor Protein Transforms Fibroblasts and Functions in Transformation by the BCR-ABL Oncogene.” Journal of Biological Chemistry 271 (38): 23255–23261.

Shah, Neil P., John M. Nicoll, Susan Branford, Timothy P. Hughes, Ronald L. Paquette, Moshe Talpaz, Claude Nicaise, Fei Huang, and Charles L. Sawyers. 2005. “Molecular Analysis of Dasatinib Resistance Mechanisms in CML Patients Identifies Novel BCR-ABL Mutations Predicted To Retain Sensitivity to Imatinib: Rationale for Combination Tyrosine Kinase Inhibitor Therapy.” Blood 106 (11): 1093–1093.

Shuai, K., J. Halpern, J. ten Hoeve, X. Rao, and C. L. Sawyers. 1996. “Constitutive Activation of STAT5 by the BCR-ABL Oncogene in Chronic Myelogenous Leukemia.” Oncogene 13 (2): 247–54.

Skorski, Tomasz, Alfonso Bellacosa, Margaret Nieborowska-Skorska, Miroslaw Majewski, Robert Martinez, John K. Choi, Rossana Trotta, et al. 1997. “Transformation of Hematopoietic Cells by BCR/ABL Requires Activation of a PI-3k/Akt-Dependent Pathway.” The EMBO Journal 16 (20): 6151–6161.

Skorski, Tomasz, P. Kanakaraj, M. Nieborowska-Skorska, M. Z. Ratajczak, SHAU-CHING Wen, G. Zon, A. M. Gewirtz, B. Perussia, and B. Calabretta. 1995. “Phosphatidylinositol-3 Kinase Activity Is Regulated by BCR/ABL and Is Required for the Growth of Philadelphia Chromosome-Positive Cells.” Blood 86 (2): 726–736.

Srivastava, Neetu, Raki Sudan, and William Garrow Kerr. 2013. “Role of Inositol Poly-Phosphatases and Their Targets in T Cell Biology.” Frontiers in Immunology 4. https://doi.org/10.3389/fimmu.2013.00288.

Steelman, L. S., S. C. Pohnert, J. G. Shelton, R. A. Franklin, F. E. Bertrand, and J. A. McCubrey. 2004. “JAK/STAT, Raf/MEK/ERK, PI3K/Akt and BCR-ABL in Cell Cycle Progression and Leukemogenesis.” Leukemia 18 (2): 189–218. https://doi.org/10.1038/sj.leu.2403241.

Taylor, Vanessa, Michelle Wong, Christian Brandts, Linda Reilly, Nicholas M. Dean, Lex M. Cowsert, Shonna Moodie, and David Stokoe. 2000. “5′ Phospholipid Phosphatase SHIP-2 Causes Protein Kinase B Inactivation and Cell Cycle Arrest in Glioblastoma Cells.” Molecular and Cellular Biology 20 (18): 6860–6871.

Trb, Thomas, Wonjae E. Choi, Gert Wolf, Elizabeth Ottinger, YunJun Chen, Michael Weiss, and Steven E. Shoelson. 1995. “Specificity of the PTB Domain of Shc for β Turn-Forming Pentapeptide Motifs Amino-Terminal to Phosphotyrosine.” Journal of Biological Chemistry 270 (31): 18205–8. https://doi.org/10.1074/jbc.270.31.18205.

Voncken, J. W., V. Kaartinen, P. K. Pattengale, W. T. Germeraad, J. Groffen, and N. Heisterkamp. 1995. “BCR/ABL P210 and P190 Cause Distinct Leukemia in Transgenic Mice.” Blood 86 (12): 4603–11.

Warmuth, Markus, Manuela Bergmann, Andrea Prie\ss, Kathrin Häuslmann, Bertold Emmerich, and Michael Hallek. 1997. “The Src Family Kinase Hck Interacts with Bcr-Abl by a Kinase-Independent Mechanism and Phosphorylates the Grb2-Binding Site of Bcr.” Journal of Biological Chemistry 272 (52): 33260–33270.

Weisberg, Ellen, Paul W. Manley, Werner Breitenstein, Josef Brüggen, Sandra W. Cowan-Jacob, Arghya Ray, Brian Huntly, et al. 2005. “Characterization of AMN107, a Selective Inhibitor of Native and Mutant Bcr-Abl.” Cancer Cell 7 (2): 129–41. https://doi.org/10.1016/j.ccr.2005.01.007.

Wisniewski, David, Annabel Strife, Steve Swendeman, Hediye Erdjument-Bromage, Scott Geromanos, W. Michael Kavanaugh, Paul Tempst, and Bayard Clarkson. 1999. “A Novel SH2-Containing Phosphatidylinositol 3, 4, 5-Trisphosphate 5-Phosphatase (SHIP2) Is Constitutively Tyrosine Phosphorylated and Associated with Src Homologous and Collagen Gene (SHC) in Chronic Myelogenous Leukemia Progenitor Cells.” Blood 93 (8): 2707–2720.

Zhao, Xun, Saghi Ghaffari, Harvey Lodish, Vladimir N. Malashkevich, and Peter S. Kim. 2002. “Structure of the Bcr-Abl Oncoprotein Oligomerization Domain.” Nature Structural Biology 9 (2): 117–20. https://doi.org/10.1038/nsb747.

Zhou, Songyang, Steven E. Shoelson, Manas Chaudhuri, Gerald Gish, Tony Pawson, Wayne G. Haser, Fred King, Tom Roberts, Sheldon Ratnofsky, and Robert J. Lechleider. 1993. “SH2 Domains Recognize Specific Phosphopeptide Sequences.” Cell 72 (5): 767–778.

Zhou, Tianjun, Lois Parillon, Feng Li, Yihan Wang, Jeff Keats, Sarah Lamore, Qihong Xu, William Shakespeare, David Dalgarno, and Xiaotian Zhu. 2007. “Crystal Structure of the T315I Mutant of Abl Kinase.” Chemical Biology & Drug Design 70 (3): 171–81. https://doi.org/10.1111/j.1747-0285.2007.00556.x.

