## Supplementary Figures for "Elucidation of protein-protein interactions necessary for maintenance of the BCR-ABL signaling complex"

Figure S1

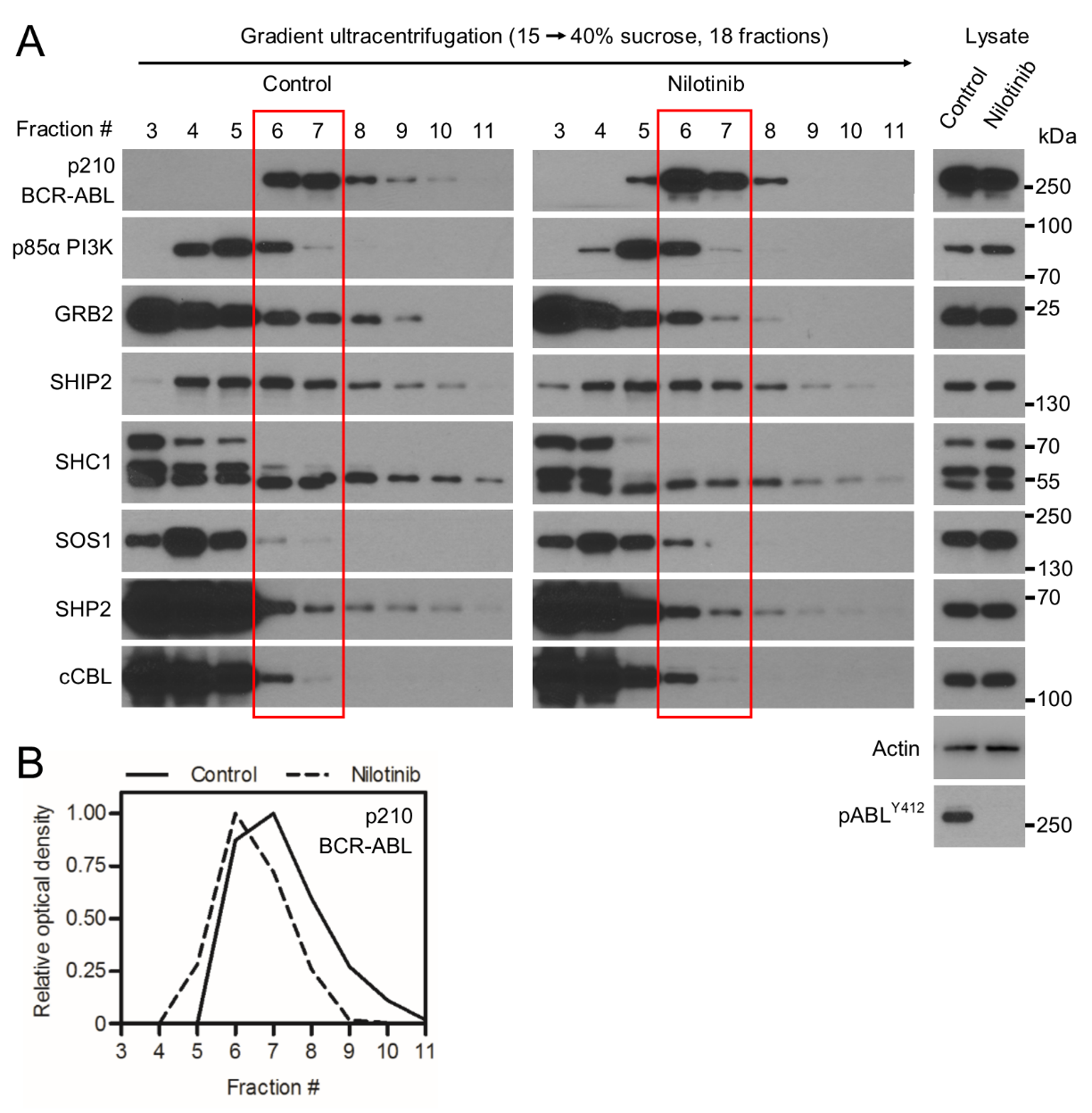

Treatment with 100 nM nilotinib produces slightly lighter BCR-ABL complexes in p210 BCR-ABL transfected 293T cells; no significant saturation of the interacting partners was found.

Figure S2

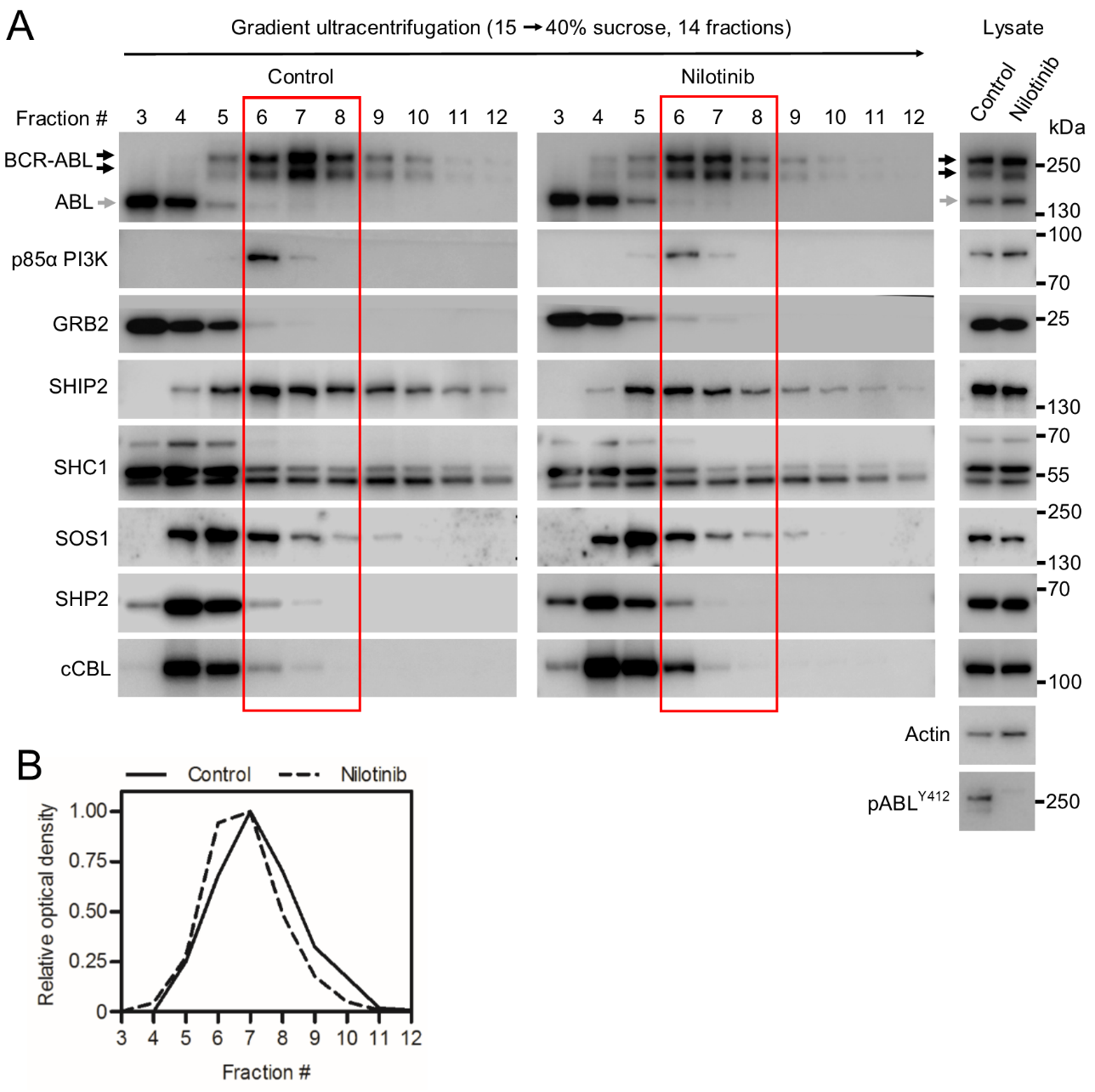

Treatment with 1 μM nilotinib does not produce a significant change in BCR-ABL complex sedimentation in BCR-ABL positive K562 cells. No significant saturation of the interacting partners was found.

Figure S3

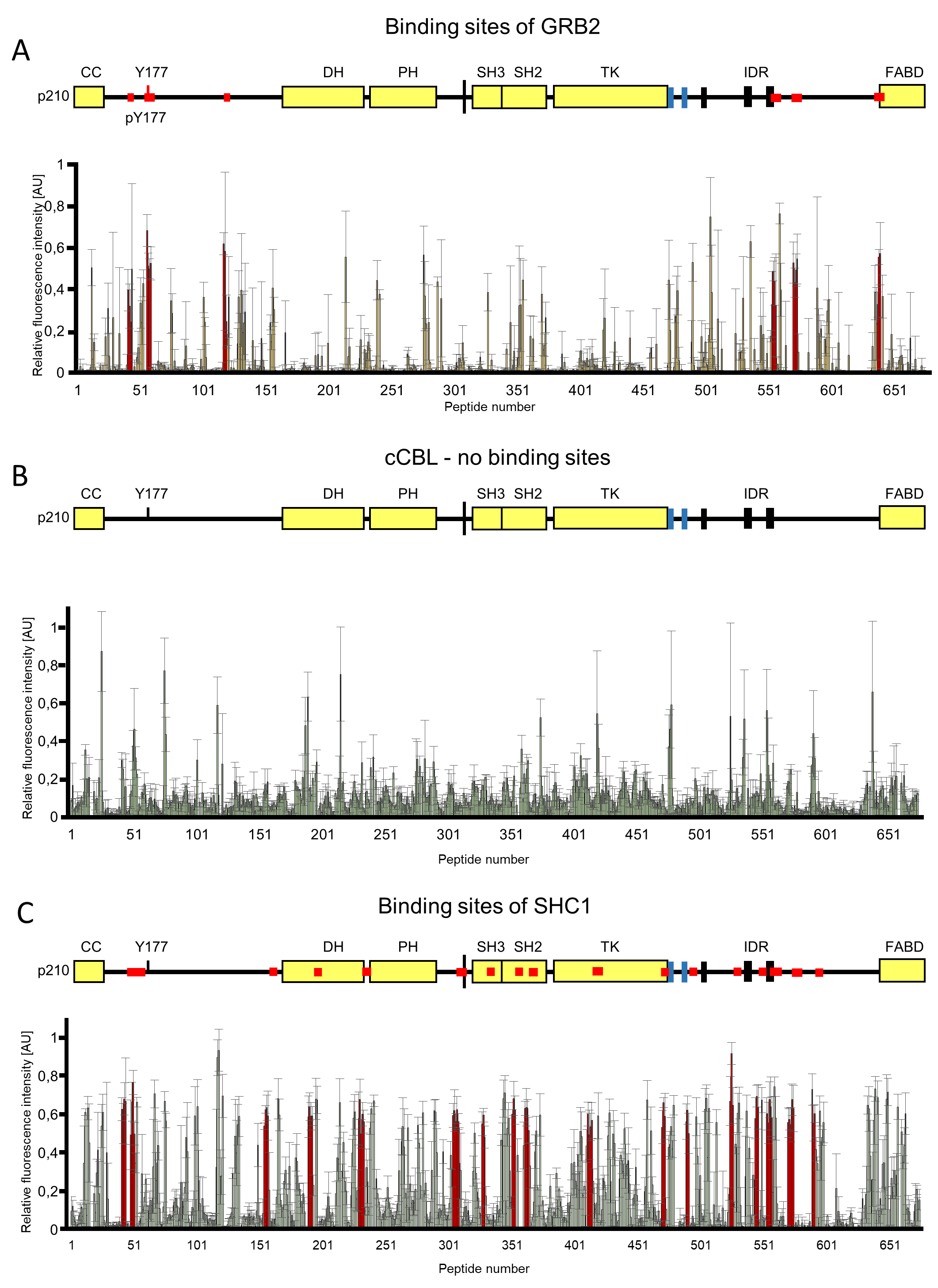

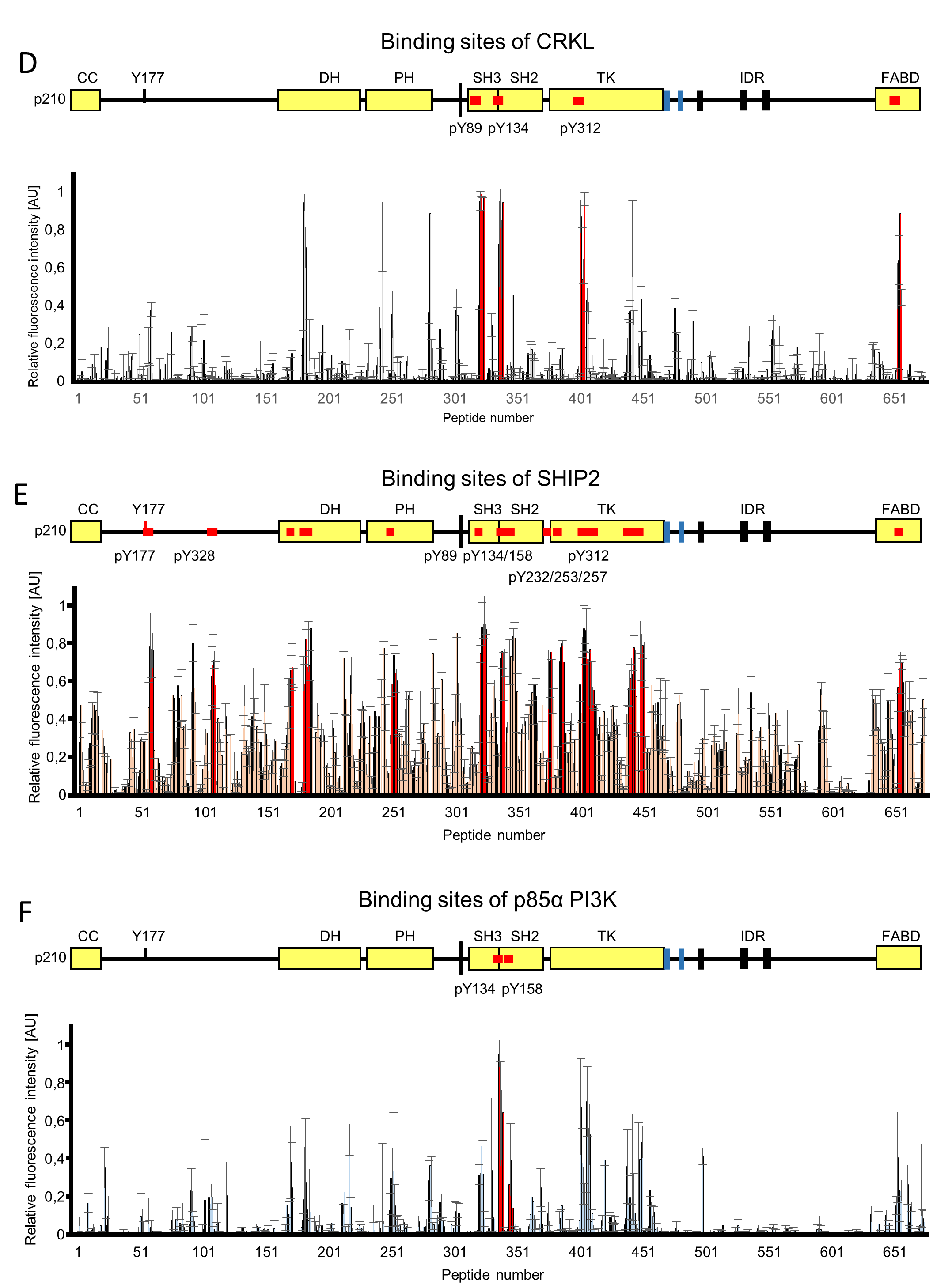

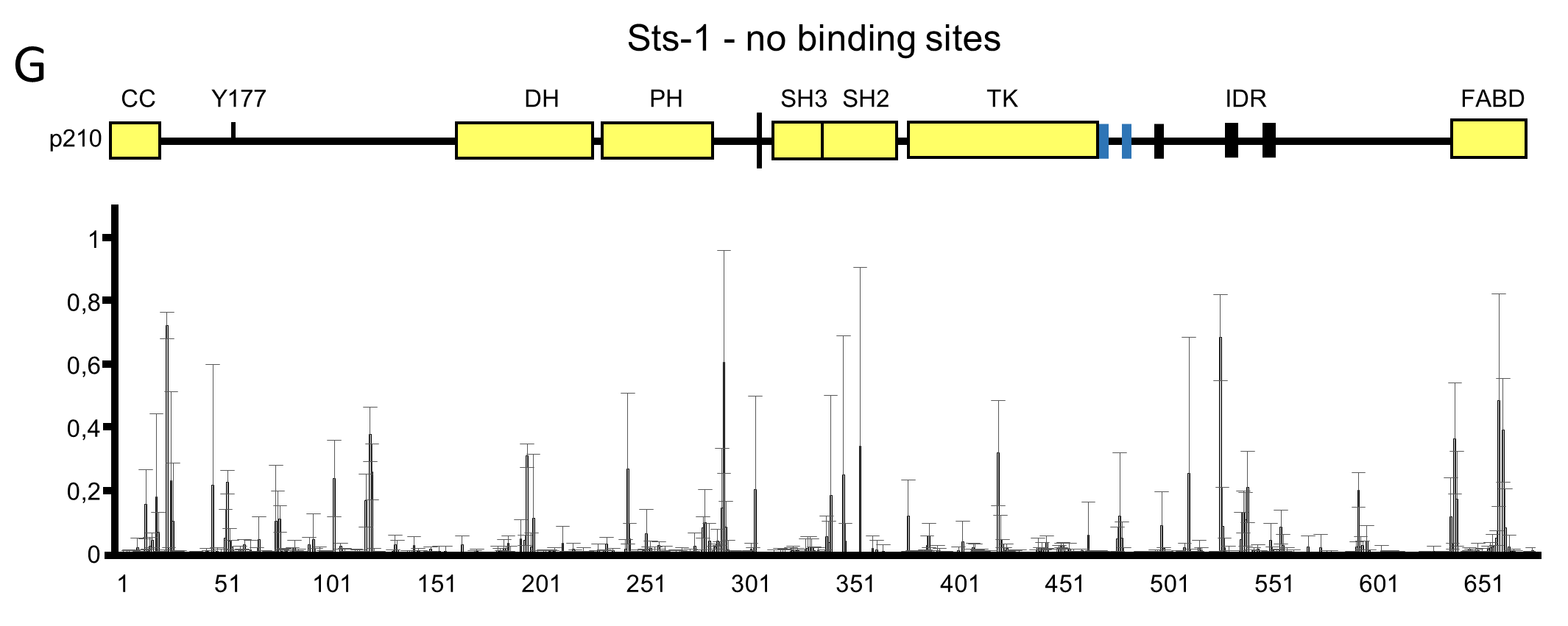
