## Supplementary Tables for "Elucidation of protein-protein interactions necessary for maintenance of the BCR-ABL signaling complex"

**Table S1** Expression vectors used in the study

| Vector | Insert | Backbone | Source/reference |
| --- | --- | --- | --- |
| BCR-ABL p210 | FLAG-BCR-ABL p210 | pcDNA3.1 | - |
| BCR-ABL p190 | FLAG-BCR-ABL p190 | pcDNA3.1 | - |
| BT-F5 | FLAG-BCR-ABL 392-2030 | pcDNA3.1 | - |
| ΔDH PH | FLAG-BCR-ABL Δ495-865 | pcDNA3.1 | - |
| ΔSH TK | FLAG-BCR-ABL Δ943-1439 | pcDNA3.1 | - |
| ΔIDR | FLAG-BCR-ABL Δ1416-1897 | pcDNA3.1 | - |
| ΔFABD2 | FLAG-BCR-ABL 1-1923 | pcDNA3.1 | - |
| KD | FLAG-BCR-ABL K271H (ABL1A n.) | pcDNA3.1 | - |
| Y177F | FLAG-BCR-ABL Y177F (BCR) | pcDNA3.1 | - |
| ΔAB | FLAG-BCR-ABL Δ190-376 | pcDNA3.1 | - |
| ΔDH | FLAG-BCR-ABL Δ468-693 | pcDNA3.1 | - |
| ΔPH1 | FLAG-BCR-ABL Δ683-826 | pcDNA3.1 | - |
| ΔSH3 | FLAG-BCR-ABL Δ935-1019 | pcDNA3.1 | - |
| ΔSH2 | FLAG-BCR-ABL Δ1026-1113 | pcDNA3.1 | - |
| ΔTK | FLAG-BCR-ABL Δ1129-1429 | pcDNA3.1 | - |
| ΔPR1 | FLAG-BCR-ABL Δ1428-1515 | pcDNA3.1 | - |
| ΔPR1-2 | FLAG-BCR-ABL Δ1416-1515 | pcDNA3.1 | - |
| ΔIDR3 | FLAG-BCR-ABL Δ1505-1669 | pcDNA3.1 | - |
| ΔPR2 | FLAG-BCR-ABL Δ1660-1878 | pcDNA3.1 | - |
| AT3 | FLAG-BCR-ABL 1-1796 | pcDNA3.1 | - |
| IDR | FLAG-BCR-ABL 1412-1919 | pcDNA3.1 | - |
| W118A | FLAG-BCR-ABL W118A (ABL1B n.) | pcDNA3.1 | - |
| S173N | FLAG-BCR-ABL S173N (ABL1B n.) | pcDNA3.1 | - |
| GRB2 | GRB2-V5-HIS | pCMV6-HIS-V5 | OriGene, modified |
| STS1 | STS1-V5-HIS | pCMV6-HIS-V5 | OriGene, modified |
| CRKL | CRKL-V5-HIS | pCMV6-HIS-V5 | OriGene, modified |
| p85α | p85a-V5-HIS | pCMV6-HIS-V5 | OriGene, modified |
| SHIP2 | SHIP2-V5-HIS | pCMV6-HIS-V5 | OriGene, modified |
| SHIP2-PD | SHIP2(P686A, D690A, R691A)-V5-HIS | pCMV6-HIS-V5 | Fafilek et al., 2018 |

**Table S2** Antibodies used in the study

| Antibody^phosphorylated motif^ | Catalog No. | Manufacturer |
| --- | --- | --- |
| Actin | sc-1615 | Santa Cruz Biotechnology |
| Actin | 3700 | Cell Signaling |
| cAbl | 2862 | Cell Signaling |
| Phospho-cAbl^Y412^ | 2865 | Cell Signaling |
| cCbl | 9794 | Cell Signaling |
| cCbl | 610441 | BD Biosciences |
| cCbl | 8447 | Cell Signaling |
| Phospho-cCbl^Y700^ | 8869 | Cell Signaling |
| Phospho-cCbl^Y731^ | 3554 | Cell Signaling |
| CrkL | 3182 | Cell Signaling |
| Phospho-CrkL^Y207^ | 3181 | Cell Signaling |
| Cy^TM^5-conjugated AffiniPure Donkey Anti-Mouse | 715-175-151 | Jackson ImmunoResearch |
| Cy^TM^5-conjugated AffiniPure Donkey Anti-Rabbit | 711-175-152 | Jackson ImmunoResearch |
| FLAG | F1804 | Sigma-Aldrich |
| GRB2 | 610111 | BD Biosciences |
| GST | G1160 | Sigma-Aldrich |
| HIS | sc-8036 | Santa Cruz Biotechnology |
| p85a-PI3K | 4257 | Cell Signaling |
| SHC1 | 610081 | BD Biosciences |
| Phospho-SHC1^Y239/240^ | 2434 | Cell Signaling |
| SHIP2 | ab166916 | Abcam |
| SHIP2 | 2730 | Cell Signaling |
| SHP2 | 610621 | BD Biosciences |
| SOS1 | sc-17793 | Santa Cruz Biotechnology |
| SOS1 | 12409 | Cell Signaling |
| STS1 | PA5-31937 | Thermo Scientific |
| V5 | R960-25 | Invitrogen |

**Table S3** GRB2 binding sites

| **Peptide number** | **Non-phosphorylated** | **phosphorylated** | **BCR-ABL residues** |
| --- | --- | --- | --- |
| 041 | SPGKARPGTARRP |  | 125-140 |
| 042 | KARPGTARRPGAA |  |  |
| 043 | PGTARRPGAAASG |  |  |
| 044 | ARRPGAAASGERD |  |  |
| **Y177_56** |  | QPGADAEKPFyVN | 170-185 |
| **Y177_57** |  | ADAEKPFyVNVEF |  |
| **Y177_58** |  | EKPFyVNVEFHHE |  |
| **Y177_59** |  | FyVNVEFHHERGL |  |
| 117 | SSRVSPSPTTYRM |  | 353-365 |
| 118 | VSPSPTTYRMFRD |  |  |
| 119 | SPTTYRMFRDKSR |  |  |
| 553 | SEKPALPRKRAGE |  | 1661-1676 |
| 554 | PALPRKRAGENRS |  |  |
| 555 | PRKRAGENRSDQV |  |  |
| 556 | RAGENRSDQVTRG |  |  |
| **570** | SSPPNLTPKPLRR |  | 1712-1727 |
| **571** | PNLTPKPLRRQVT |  |  |
| **572** | TPKPLRRQVTVAP |  |  |
| **573** | PLRRQVTVAPASG |  |  |
| **637** | RVSLRKTRQPPER |  | 1913-1925 |
| **638** | LRKTRQPPERIAS |  |  |
| **639** | TRQPPERIASGAI |  |  |

**Table S4** SHC1 binding sites

| **Peptide number** | **Non-phosphorylated** | **phosphorylated** | **BCR-ABL residues** |
| --- | --- | --- | --- |
| 041 | SPGKARPGTARRP |  | 125-140 |
| 042 | KARPGTARRPGAA |  |  |
| 043 | PGTARRPGAAASG |  |  |
| 044 | ARRPGAAASGERD |  |  |
| 048 | DDRGPPASVAALR |  | 146-161 |
| 049 | GPPASVAALRSNF |  |  |
| 050 | ASVAALRSNFERI |  |  |
| 051 | AALRSNFERIRKG |  |  |
| **154** | SSSPHLSSKGRGS |  | 464-479 |
| **155** | PHLSSKGRGSRDA |  |  |
| **156** | SSKGRGSRDALVS |  |  |
| **157** | GRGSRDALVSGAL |  |  |
| 189 | PRVQQWSHQQRVG |  | 569-584 |
| 190 | QQWSHQQRVGDLF |  |  |
| 191 | SHQQRVGDLFQKL |  |  |
| 192 | QRVGDLFQKLASQ |  |  |
| 229 | SSINEEITPRRQS |  | 689-707 |
| 230 | NEEITPRRQSMTV |  |  |
| 231 | ITPRRQSMTVKKG |  |  |
| 232 | RRQSMTVKKGEHR |  |  |
| 233 | SMTVKKGEHRQLL |  |  |
| 304 | GFLNVIVHSATGF |  | 914-932 |
| 305 | NVIVHSATGFKQS |  |  |
| 306 | VHSATGFKQSSKA |  |  |
| 307 | ATGFKQSSKALQR |  |  |
| 308 | FKQSSKALQRPVA |  |  |
| 327 | TLSITKGEKLRVL |  | 983-995 |
| 328 | ITKGEKLRVLGYN |  |  |
| 329 | GEKLRVLGYNHNG |  |  |
| **351** | VRESESSPGQRSI |  | 1055-1067 |
| **352** | SESSPGQRSISLR |  |  |
| **353** | SPGQRSISLRYEG |  |  |
| **361** | DGKLYVSSESRFN |  | 1085-1100 |
| **362** | LYVSSESRFNTLA |  |  |
| **363** | SSESRFNTLAELV |  |  |
| **364** | SRFNTLAELVHHH |  |  |
| 411 | NRQEVNAVVLLYM |  | 1235-1250 |
| 412 | EVNAVVLLYMATQ |  |  |
| 413 | AVVLLYMATQISS |  |  |
| 414 | LLYMATQISSAME |  |  |
| **470** | KELGKQGVRGAVS |  | 1412-1424 |
| **471** | GKQGVRGAVSTLL |  |  |
| **472** | GVRGAVSTLLQAP |  |  |
| 489 | EPAVSPLLPRKER |  | 1469-1481 |
| 490 | VSPLLPRKERGPP |  |  |
| 491 | LLPRKERGPPEGG |  |  |
| **524** | NGALRESGGSGFR |  | 1574-1586 |
| **525** | LRESGGSGFRSPH |  |  |
| **526** | SGGSGFRSPHLWK |  |  |
| **544** | EWRSVTLPRDLQS |  | 1634-1646 |
| **545** | SVTLPRDLQSTGR |  |  |
| **546** | LPRDLQSTGRQFD |  |  |
| 553 | SEKPALPRKRAGE |  | 1661-1679 |
| 554 | PALPRKRAGENRS |  |  |
| 555 | PRKRAGENRSDQV |  |  |
| 556 | RAGENRSDQVTRG |  |  |
| 557 | ENRSDQVTRGTVT |  |  |
| **570** | SSPPNLTPKPLRR |  | 1712-1730 |
| **571** | PNLTPKPLRRQVT |  |  |
| **572** | TPKPLRRQVTVAP |  |  |
| **573** | PLRRQVTVAPASG |  |  |
| **574** | RQVTVAPASGLPH |  |  |
| **589** | TSKGPAEESRVRR |  | 1769-1781 |
| **590** | GPAEESRVRRHKH |  |  |
| **591** | EESRVRRHKHSSE |  |  |

**Table S5** CRKL binding sites

| **Peptide number** | **Non-phosphorylated** | **phosphorylated** | **BCR-ABL residues** |
| --- | --- | --- | --- |
| Y89 - ABL_320 |  | PSENDPNLFVALy | 959-977 |
| Y89 - ABL_321 |  | NDPNLFVALyDFV |  |
| Y89 - ABL_322 |  | NLFVALyDFVASG |  |
| Y89 - ABL_323 |  | VALyDFVASGDNT |  |
| Y89 - ABL_324 |  | yDFVASGDNTLSI |  |
| Y134 (115)_336 |  | NGQGWVPSNyITP | 1010-1025 |
| Y134 (115)_337 |  | GWVPSNyITPVNS |  |
| Y134 (115)_338 |  | PSNyITPVNSLEK |  |
| Y134 (115)_339 |  | yITPVNSLEKHSW |  |
| Y312_401 |  | LLGVCTREPPFyI | 1205-1220 |
| Y312_402 |  | VCTREPPFyIITE |  |
| Y312_403 |  | REPPFyIITEFMT |  |
| Y312_404 |  | PFyIITEFMTYGN |  |
| 653 | LEAGKNLYTFCVS |  | 1961-1976 |
| 654 | GKNLYTFCVSYVD |  |  |
| 655 | LYTFCVSYVDSIQ |  |  |
| 656 | FCVSYVDSIQQMR |  |  |

**Table S6** SHIP2 binding sites

| **Peptide number** | **Non-phosphorylated** | **phosphorylated** | **BCR-ABL residues** |
| --- | --- | --- | --- |
| 057 | ADAEKPFYVNVEF | ADAEKPFyVNVEF | 173-185 |
| 058 | EKPFYVNVEFHHE | EKPFyVNVEFHHE |  |
| 059 | FYVNVEFHHERGL | FyVNVEFHHERGL |  |
| Y328_106 |  | SPRSFEDCGGGyT | 320-335 |
| Y328_107 |  | SFEDCGGGyTPDC |  |
| Y328_108 |  | DCGGGyTPDCSSN |  |
| Y328_109 |  | GGyTPDCSSNENL |  |
| 169 | ILASEETYLSHLE |  | 509-521 |
| 170 | SEETYLSHLEALL |  |  |
| 171 | TYLSHLEALLLPM |  |  |
| 179 | PVLTSQQIETIFF |  | 539-564 |
| 180 | TSQQIETIFFKVP |  |  |
| 181 | QIETIFFKVPELY |  |  |
| 182 | TIFFKVPELYEIH |  |  |
| 183 | FKVPELYEIHKEF |  |  |
| 184 | PELYEIHKEFYDG |  |  |
| 185 | YEIHKEFYDGLFP |  |  |
| 249 | GKTQQYDCKWYIP |  | 749-767 |
| 250 | QQYDCKWYIPLTD |  |  |
| 251 | DCKWYIPLTDLSF |  |  |
| 252 | WYIPLTDLSFQMV |  |  |
| 253 | PLTDLSFQMVDEL |  |  |
| 320 | PSENDPNLFVALY | PSENDPNLFVALy | 962-980 |
| 321 | NDPNLFVALYDFV | NDPNLFVALyDFV |  |
| 322 | NLFVALYDFVASG | NLFVALyDFVASG |  |
| 323 | VALYDFVASGDNT | VALyDFVASGDNT |  |
| 324 | YDFVASGDNTLSI | yDFVASGDNTLSI |  |
| Y134 (115)_336 |  | NGQGWVPSNyITP | 1010-1025 |
| Y134 (115)_337 |  | GWVPSNyITPVNS |  |
| Y134 (115)_338 |  | PSNyITPVNSLEK |  |
| Y134 (115)_339 |  | yITPVNSLEKHSW |  |
| Y232 (251)_374 |  | RNKPTVYGVSPNy | 1124-1136 |
| Y232 (251)_375 |  | PTVYGVSPNyDKW |  |
| Y232 (251)_376 |  | YGVSPNyDKWEME |  |
| Y253_382 |  | MKHKLGGGQyGEV | 1148-1163 |
| Y253_383 |  | KLGGGQyGEVYEG |  |
| Y253_384 |  | GGQyGEVYEGVWK |  |
| Y253_385 |  | yGEVYEGVWKKYS |  |
| Y257_383 |  | KLGGGQYGEVyEG | 1150-1165 |
| Y257_384 |  | GGQYGEVyEGVWK |  |
| Y257_385 |  | YGEVyEGVWKKYS |  |
| Y257_386 |  | VyEGVWKKYSLTV |  |
| Y312_401 |  | LLGVCTREPPFyI | 1205-1220 |
| Y312_402 |  | VCTREPPFyIITE |  |
| Y312_403 |  | REPPFyIITEFMT |  |
| Y312_404 |  | PFyIITEFMTYGN |  |
| 405 | IITEFMTYGNLLD |  | 1217-1238 |
| 406 | EFMTYGNLLDYLR |  |  |
| 407 | TYGNLLDYLRECN |  |  |
| 408 | NLLDYLRECNRQE |  |  |
| 409 | DYLRECNRQEVNA |  |  |
| 410 | RECNRQEVNAVVL |  |  |
| 438 | AYNKFSIKSDVWA |  | 1316-1337 |
| 439 | KFSIKSDVWAFGV |  |  |
| 440 | IKSDVWAFGVLLW |  |  |
| 441 | DVWAFGVLLWEIA |  |  |
| 442 | AFGVLLWEIATYG |  |  |
| 443 | VLLWEIATYGMSP |  |  |
| 447 | PYPGIDLSQVYEL |  | 1343-1358 |
| 448 | GIDLSQVYELLEK |  |  |
| 449 | LSQVYELLEKDYR |  |  |
| 450 | VYELLEKDYRMER |  |  |
| 652 | SAVLEAGKNLYTF |  | 1958-1976 |
| 653 | LEAGKNLYTFCVS |  |  |
| 654 | GKNLYTFCVSYVD |  |  |
| 655 | LYTFCVSYVDSIQ |  |  |
| 656 | FCVSYVDSIQQMR |  |  |

**Table S7** p85a-PI3K binding sites

| **Peptide number** | **Non-phosphorylated** | **phosphorylated** | **BCR-ABL residues** |
| --- | --- | --- | --- |
| Y134 (115)_Peptide_336 |  | NGQGWVPSNyITP | 1008-1022 |
| Y134 (115)_Peptide_337 |  | GWVPSNyITPVNS |  |
| Y134 (115)_Peptide_338 |  | PSNyITPVNSLEK |  |
| Y134 (115)_Peptide_339 |  | yITPVNSLEKHSW |  |
| Y158 (139)_Peptide_343 |  | WYHGPVSRNAAEy | 1030-1046 |
| Y158 (139)_Peptide_344 |  | GPVSRNAAEyLLS |  |
| Y158 (139)_Peptide_345 |  | SRNAAEyLLSSGI |  |
| Y158 (139)_Peptide_346 |  | AAEyLLSSGINGS |  |
